# Spatial patterns of diversity in forest birds of peninsular India

**DOI:** 10.1101/2025.04.28.650994

**Authors:** Naman Goyal, Archita Sharma, Vishwa Jagati, Akshay Herur, Arunima Jain, Abhishek Gopal, Jahnavi Joshi, VV Robin

## Abstract

Biodiversity is structured via complex interactions among ecological, geological, and climatic processes. Regions with high heterogeneity in climate and topography are known to harbor fine scale patterns in diversity which are often overlooked in global scale analyses. Here, we investigate spatial phylogenetic patterns of forest bird diversity across peninsular India, a region with high topographic and climatic heterogeneity. Using a comprehensive global bird phylogeny, we employ metrics such as Phylogenetic Diversity (PD), Phylogenetic Endemism (PE), and their relative forms (RPD, RPE) to quantify evolutionary history and endemism of forest birds in peninsular India. We examine the roles of contemporary climate, historic climatic stability, and topography in shaping these patterns. Our results reveal a strong gradient in diversity, with the southern Western Ghats acting as a major hotspot for both taxonomic and phylogenetic diversity and endemism. We detect distinct ecological and phylogenetic community structures across the peninsula, likely shaped by regional species pools and biogeographic barriers. Areas with greater topographic complexity, higher precipitation, and greater historic climatic stability were found to support high diversity. Our study provides critical insights for biogeography in this understudied yet highly biodiverse region for forest birds.

## Introduction

Biodiversity is unevenly distributed across the globe and there has been much interest to understand the processes that drive these patterns. With unprecedented declines in biodiversity (Pereira et al. 2010), we are not only losing species, but also areas with distinct evolutionary history (Veron et al. 2017) and ecosystem services (Dobson et al. 2006). Hence, understanding the drivers of different facets of diversity is crucial for planning better conservation strategies.

Geological, ecological, and environmental processes are known to interactively structure patterns in diversity across space and time (Wiens and Donoghue 2004). Some of the key hypotheses explaining biodiversity patterns focus on topographic complexity, energy availability, and historical climatic stability. Mountains, for example, harbor higher-than-expected biodiversity due to steep climatic gradients over short distances and their role as barriers to dispersal, which promote isolation and species diversification (Rahbek, Borregaard, Antonelli, et al. 2019). Similarly, regions with high energy availability are expected to support greater diversity by providing the resources necessary for species coexistence (Araújo et al. 2008; Evans, Warren, and Gaston 2005; Hawkins et al. 2003). Historical climatic stability is also a strong predictor of diversity, as reduced climatic fluctuations over evolutionary timescales allow for lineage persistence and accumulation (Araújo et al. 2008; Carnaval et al. 2009; Gómez et al. 2007; Wiens and Donoghue 2004). While these broad-scale mechanisms are well-recognized, understanding how they interact to influence regional patterns of biodiversity—particularly in topographically and climatically complex landmasses—remains a critical challenge.

Traditional methods of quantifying biodiversity and identifying conservation hotspots have largely relied on species richness (SR) (Myers et al. 2000). However, more recent approaches increasingly incorporate phylogenetic information to assess biodiversity patterns in an evolutionary context (Rosauer et al. 2009; Nitta et al. 2023; Paúl et al. 2023). These measures enable the assessment of evolutionary uniqueness in communities and the identification of regions with high evolutionary significance. Furthermore, they elucidate the influence of historical processes by revealing patterns in lineage diversification, extinction, and persistence (Rosauer et al. 2009; Nitta et al. 2023; Paúl et al. 2023). Phylogenetic Diversity (PD) and Phylogenetic Endemism (PE) are among the most widely used metrics for evaluating biodiversity through a phylogenetic lens. PD quantifies the total branch length of a phylogeny connecting the species in an area (Faith 1992), while PE weights those branch lengths by the relative geographic ranges of the species involved (Rosauer et al. 2009). Both metrics incorporate evolutionary history and provide valuable insights into how long-term ecological and geological processes have shaped current biodiversity patterns.

Peninsular India is a region with complex geoclimatic history and has high heterogeneity in topography and climate (Karanth 2017; Joshi and Karanth 2011). It consists of major topographical features such as the Deccan plateau, the Western Ghats mountains, the Eastern Ghats mountains and the Chota Nagpur plateau (Figure 1). Furthermore based on Koppen climate classification, peninsular India consists of climatic zones ranging from tropical wet to arid climatic conditions (Beck et al. 2023; Jain et al. 2022).

**Figure 1.**
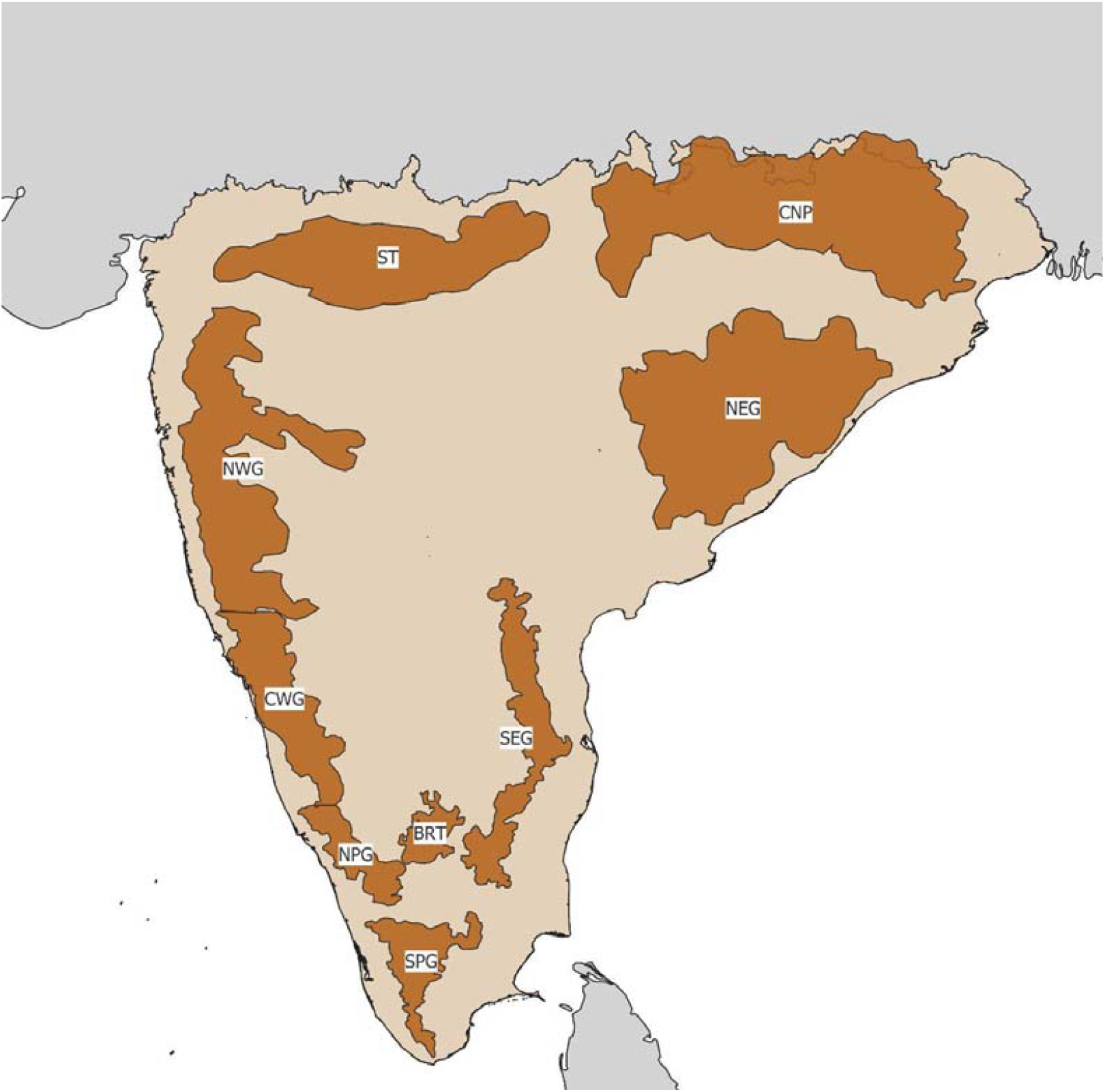
Map of peninsular India depicting major topographic zones (>800m elevation) - the Western Ghats, divided into northern (NWG), central (CWG), and southern regions (NPG and SPG), the Eastern Ghats divided into north (NEG) and south (SEG), the Satpuras (ST), the Chota Nagpur Plateau (CNP), and the Biligirirangan Hills (BRT).

Patterns in distribution of biodiversity have been well documented in the Western Ghats (WG), a global biodiversity hotspot in peninsular India. Studies on diversity patterns of several taxa show the presence of a latitudinal diversity gradient in the WG (Gopal et al. 2023; Bharti et al. 2021; Bose et al. 2019; Divya, Ramesh, and Karanth 2021; Page and Shanker 2020; Davidar, Puyravaud, and Leigh 2005; Daniels 1992). Moreover, patterns in endemism for plants (Bose et al. 2019; Gopal et al. 2023), snails (Aravind, Rajashekhar, and Madhyastha 2005), centipedes (Bharti et al. 2021), and amphibians (Daniels 1992) show that southern WG has a higher proportion of endemic species compared to its northern stretches. Further, The presence of old lineages of centipedes (Bharti et al. 2021), and both old and young lineages of plants (Gopal et al. 2023) supports the idea that southern WG has been a historic climatic refuge, acting both as a museum and cradle of diversity (Joshi and Karanth 2013). Environmental variation, topographic complexity and historic climatic stability have played a role in driving these patterns in the WG (Page and Shanker 2020; Gopal et al. 2023; Bharti et al. 2021; Divya, Ramesh, and Karanth 2021; Bose et al. 2019).

Our understanding of these patterns are limited at a broader peninsular India scale, with the inclusion of other mountain ranges such as the Eastern Ghats. (Ramachandran et al. 2017) examined the distribution of birds at a subspecies level in peninsular India, and found that several distribution boundaries coincided with physical or climatic divides within the peninsula, suggesting a role of both climate and topography in driving subspecies level distributional patterns. In addition, they find that wet regions that have been historically stable – potential refugia – harbor a higher proportion of endemism for birds. However, they relied only on subspecies level natural history information and did not incorporate phylogenetic information in assessing spatial patterns in diversity patterns for birds in peninsular India.

In this study, we investigate spatial phylogenetic patterns in the distribution of forest birds across peninsular India using the latest global bird phylogeny (McTavish et al. 2024). Specifically, we test the roles of climate and topography in shaping these patterns. To do so, we employ a suite of phylogenetic metrics, Phylogenetic Diversity (PD), Phylogenetic Endemism (PE), Relative Phylogenetic Diversity (RPD), and Relative Phylogenetic Endemism (RPE), which allow us to assess different aspects of evolutionary history and endemism within communities. While PD and PE help quantify the total evolutionary history and the geographic restriction of that history in a given region, RPD and RPE compare observed patterns to a null expectation (Mishler et al. 2014), thus highlighting regions with disproportionately old or young lineages. These metrics together enable us to identify areas acting as evolutionary hotspots, detect potential refugia, and understand how historical processes have contributed to current biodiversity patterns in peninsular India, including less-studied regions like the Eastern Ghats.

In particular, we focus on the following set of questions with specific expectations:

1. Do lower latitudes show greater diversity compared to the rest of peninsular India? Based on the latitudinal diversity gradient hypothesis, we expect higher diversity in regions in the lower latitudes, such as the southern Western Ghats, compared to other regions in peninsular India.
2. Do different regions within the peninsula show differences in their community structure in an ecological and phylogenetic perspective? Given that local community assemblages could be shaped by regional species pools (Cornell and Harrison 2014), we expect to find clusters of taxonomically and phylogenetically similar communities across the landscape. Identifying these patterns will provide insights into biogeographic regionalization within peninsular India using both taxonomic and phylogenetic information for species.
3. Does historic climatic stability, contemporary climate and topography predict diversity across peninsular India? Historic climatic stability, contemporary climate and topography are known to drive patterns in diversity both at global scales and regional scales (Araújo et al. 2008; Gómez et al. 2007; Carnaval et al. 2009; Evans, Warren, and Gaston 2005; Rahbek, Borregaard, Antonelli, et al. 2019). We examine the roles of historic climate stability, contemporary climate, and topography in driving the patterns in diversity for forest birds in peninsular India. We expect that areas with high topographical complexity, high precipitation (wetter areas), and high historic climatic stability will have higher diversity of forest birds.

## Methods

### Study Region and Species Selection

We define peninsular India as the inverted triangle part of the Indian subcontinent, bound by the Western Ghats in the west, the Eastern Ghats in the east, and the Satpuras and Chota Nagpur plateau in the north (Figure 1). There are approximately 600 species of birds that are known to occur in this region, with a small variation in the number depending on the source of taxonomy. We further restricted our analyses only to the resident, breeding, terrestrial, forest bird species.

### Species distribution and phylogenetic data

We used the distribution maps for birds of the world from Birdlife International (BirdLife International, Handbook of the Birds of the World 2023) clipped to the extent of our study area (QGIS v 3.24.3; (QGIS Development Team 2024)). Furthermore, we filtered the dataset to include only birds that are native resident breeding species (values 1 and 2 under Birdlife metadata “Season”, and values 1 under “Origin”), and omitted migratory species that breed elsewhere from downstream analyses, resulting in 329 species. We used the AVONET dataset (Tobias et al. 2022) to classify the species based on the predominant habitat they use. Since the focus of this study was only on forest and woodland birds, we filtered the dataset to retain only species that predominantly use these habitats including Riverine habitats. Including species from non-forest habitats would dilute the ecological signal and reduce the interpretability of the results. Filtering by forest and woodland habitat resulted in a total of 188 species. We further renamed species following Clements/eBird 2023 taxonomy for downstream analyses to be able to use the latest bird phylogeny (McTavish et al. 2024). We pruned the phylogenetic tree to retain only the 188 species for downstream analyses (Figure S1). We also extracted the occurrence records for the final set of 188 species from eBird dataset using the package *Auk* in R.

### Estimating Avian diversity metrics

To estimate the diversity metrics, we first generated presence-absence matrices at 3 scales - 10km, 25km and 50km. We chose these three relatively coarse scales as BirdLife range maps that are based on expert opinion and may not represent fine scale species distributions (Warudkar et al. 2022). The presence-absence matrices were generated (*lets.presab.grids* in the package LetsR (Vilela and Villalobos 2015)) and we estimated the Species Richness (SR) as the sum of all species present in each grid. The weighted endemism (WE) was estimated as the sum of presence of all the species occurring in a grid by the number of grids occupied by each species in the region of interest. We estimated Phylogenetic Diversity (PD), Phylogenetic Endemism (PE), Relative Phylogenetic Diversity (RPD), and Relative Phylogenetic Endemism (RPE) (*rand_test_cpr* of package *canaper* (Nitta et al. 2023)) simultaneously with testing for null models. For generating the null models we used the “swap” algorithm of the package *vegan (Oksanen et al. 2025)*. Briefly, this algorithm swaps 2×2 submatrices to randomize the community matrix while maintaining the row and column sums (Gotelli and Entsminger 2003). To ensure that we have randomized our null models sufficiently, we ran optimization for *n_itr* and *n_reps* as per the recommendations (Nitta et al. 2023) (Figure S2). Based on our optimization results, we found that 200,000 n_reps and 1,000 n_itrs were enough to randomize a community at a 25 x 25 km scale (Figure S2 and S3). Whereas, n_reps had to be increased to 2,000,000 at a scale of 10 x 10 km to get a sufficiently randomized null community (Figure S3). We also estimated time-integrated lineage diversity (TILD) using a custom R script. TILD is a deep-time complement to PD, it is estimated by integrating the area under a lineage-through time plot where the number of lineages are log-transformed (Dexter, Segovia, and Griffiths 2019). Further, to identify different types of endemism centres, we used a two step process called Categorical Analysis of Neo and Paleo Endemism (CANAPE) implemented in package *canaper*. Briefly, this algorithm identifies grid cells with significantly high values (one-tailed test, α = 0.05) in the numerator, denominator, or both components of RPE, indicating elevated endemism. Among these, cells are then classified based on the RPE ratio (two-tailed test, α = 0.05) into three categories: (1) paleo endemism (high RPE), (2) neo endemism (low RPE), and (3) mixed endemism (high in both numerator and denominator, but non-significant RPE ratio). Mixed endemism indicates a combination of rare long and short branches. Grid cells with high significance (α = 0.01) in both numerator and denominator were further designated as super endemism sites (Mishler et al. 2014; Nitta et al. 2023).

### Assessing the biotic turnover in forest birds across peninsular India

To assess the structure of communities across peninsular India for the selected species, we used a Grade of Membership model to cluster communities based on presence-absence data, as implemented in the R package *Ecostructure* (White et al. 2019, 2021). Grade of Membership models allow units of analyses, such as geographic locations or species, to belong to multiple clusters (White et al. 2019, 2021). We ran this model for values of K from 2 to 9 with 10,000 iterations for each K, and selected the best model based on estimated log likelihood for each K. We plotted the membership proportions using pie charts for each grid using the *ggplot2* package in R (Wickham 2016). Further, for each K, we identified the top species contributing to each cluster using the function *ExtractTopFeatures* in the *ecostructure* package (White et al. 2019, 2021) (Table S2). To evaluate the optimal value of K for peninsular India, we plotted the log likelihood values for each K from 2 - 9, and identified the optimal value of K where the increase in log likelihood starts flattening, that is the elbow of the plot. This method is used regularly to estimate the optimal value of K in structure like methods in population genomics (Evanno, Regnaut, and Goudet 2005).

Further, to assess the turnover in phylogenetic diversity across peninsular India, we estimated the phylogenetic beta diversity using Sorenson’s Phylogenetic Index (Leprieur et al. 2012). We used the function *phylosor* from package *picante* in R to generate a distance matrix of pairwise similarity between sites (Kembel et al. 2010). Further, we carried out a K-means clustering using the distance matrix for values of K from 1 to 10 to identify regions of similar phylogenetic diversity. We calculated the total within-cluster sum of squares (WSS) to evaluate the appropriate value for K. The WSS represents the sum of the squared distances between each data point and its assigned cluster centroid. We then plotted the WSS values against the corresponding K values, creating an elbow plot. The ‘elbow’ point in the plot, where the rate of decrease in WSS sharply diminishes, was identified as the optimal K.

### Environmental Correlates of Avian Diversity

We used a set of climatic and topographic variables that capture the spatial and temporal heterogeneity in the region and are known to influence patterns in distribution of birds (Acevedo and Sandel 2021; Ramachandran et al. 2017). The data were available at 1km x 1km resolution, and aggregated to 25km x 25km resolution. We checked for autocorrelation for all the variables using the *corrplot* package in R and selected variables which had Pearson’s correlation value of less than 0.5 or more than −0.5. In total we selected 12 climatic and topographic variables (Table 1).

**Table 1.** List of climatic and topographic variables used in this study to identify the drivers of species richness, phylogenetic diversity and phylogenetic endemism in forest birds of peninsular India.

| Type | Variable | Description | Source |
| --- | --- | --- | --- |
| Climatic | Bio9 | Mean Temperature of Driest Quarter | (Karger et al. 2017) |
|  | Bio10 | Mean Temperature of Warmest Quarter | (Karger et al. 2017) |
|  | Bio11 | Mean Temperature of Coldest Quarter | (Karger et al. 2017) |
|  | Bio16 | Precipitation of Wettest Quarter | (Karger et al. 2017) |
|  | Bio17 | Precipitation of Driest Quarter | (Karger et al. 2017) |
|  | Bio18 | Precipitation of Warmest Quarter | (Karger et al. 2017) |
|  | TCC mean | Mean of Total Cloud Cover | (Wilson and Jetz 2016) |
|  | TCC sd | Standard Deviation of Total Cloud Cover | (Wilson and Jetz 2016) |
|  | LGM_CCV | Quaternary Climate Change Velocity | This study |
| Topographic | Aspect cosine | Cosine of Aspect | (Amatulli et al. 2018) |
|  | Aspect sine | Sine of Aspect | (Amatulli et al. 2018) |
|  | Elevation | Elevation relative to mean sea level | (Amatulli et al. 2018) |
|  | TRI | Topographic Ruggedness Index | (Amatulli et al. 2018) |

To estimate the climatic stability of the region since the last glacial maxima (LGM) we relied on two main variables – the Annual Mean Temperature (Bio1) and Annual Precipitation (Bio12). We obtained the data at 1km x 1km resolution at 100-year time intervals from the Trace21k dataset (Karger et al. 2023). We used climate change velocity to measure the climatic stability of the study area since the LGM. Climate change velocity is a measure of local climatic displacement over an area and is calculated by dividing the change in climate over time by change in climate over space for a given area (Loarie et al. 2009). Climate change velocity is a measure of the rate at which the species would have to shift their ranges to track changes in climate (Loarie et al. 2009; Sandel et al. 2011). The data was first aggregated to a 25km x 25km scale in R using the package *raster*. We then used the package *VoCC* in R to estimate the gradient-based climate change velocity (García Molinos et al. 2019). We first estimated the climate change velocity for Bio1 and Bio12 separately, then rescaled the velocities to a scale of 0 to 1 by normalizing the resultant rasters. Here, 0 means lowest velocity or a climatically stable region and 1 means highest velocity of climate change or climatically unstable region. We finally combined both temperature and precipitation climate change velocities into a single raster by adding the layers together. We checked for correlation between climate change velocity with the 12 selected variables using the same method detailed above.

We assessed the role of climatic and topographic variables at 25 x 25 km scale. First, we scaled all the predictor variables and ran linear regression models for Species Richness (SR), Phylogenetic Diversity (PD), and Phylogenetic Endemism (PE). Second, we ran 3 sets of models for each response variable, 1) with only climatic variables, 2) with only topographic variables, and 3) combined climatic and topographic variables. We tested for spatial independence in residuals for all the linear regression models using the function *correlog* of the package *pgirmess* in R to estimate the Moran’s I statistic at different distance classes (Giraudoux 2013). Since we found significant spatial autocorrelation in the residuals (p < 0.05), we used spatial autoregressive models (Kissling and Carl 2008). For each model we identified all neighbors up to the first non-significant value in our correlog plots using the function *dnearneigh* in the *spdep* package in R (Bivand and Piras 2015). We then generated a spatial weights matrix using the function nb2listw using a standard coding scheme (Kissling and Carl 2008). We tested multiple spatial models in a model selection framework implemented in the *spdep* package in R. The models were then ranked according to their model performance using Akaike Information Criterion (AIC) scores (Table S3).

## Results

### Patterns in Species Richness, Phylogenetic Diversity and Phylogenetic Endemism

Our results indicate that the highest species richness (SR) and weighted endemism (WE) was found to be within the southern Western Ghats (Figure 2A and 2B). Overall, we observe a pattern consistent with latitudinal diversity gradient, where lower latitudes have higher species richness of forest birds than the higher latitudes in the Western Ghats. However, in the case of the Eastern Ghats, the pattern is reversed, where higher species richness is observed in high latitudes (Figure 2A).

**Figure 2.**
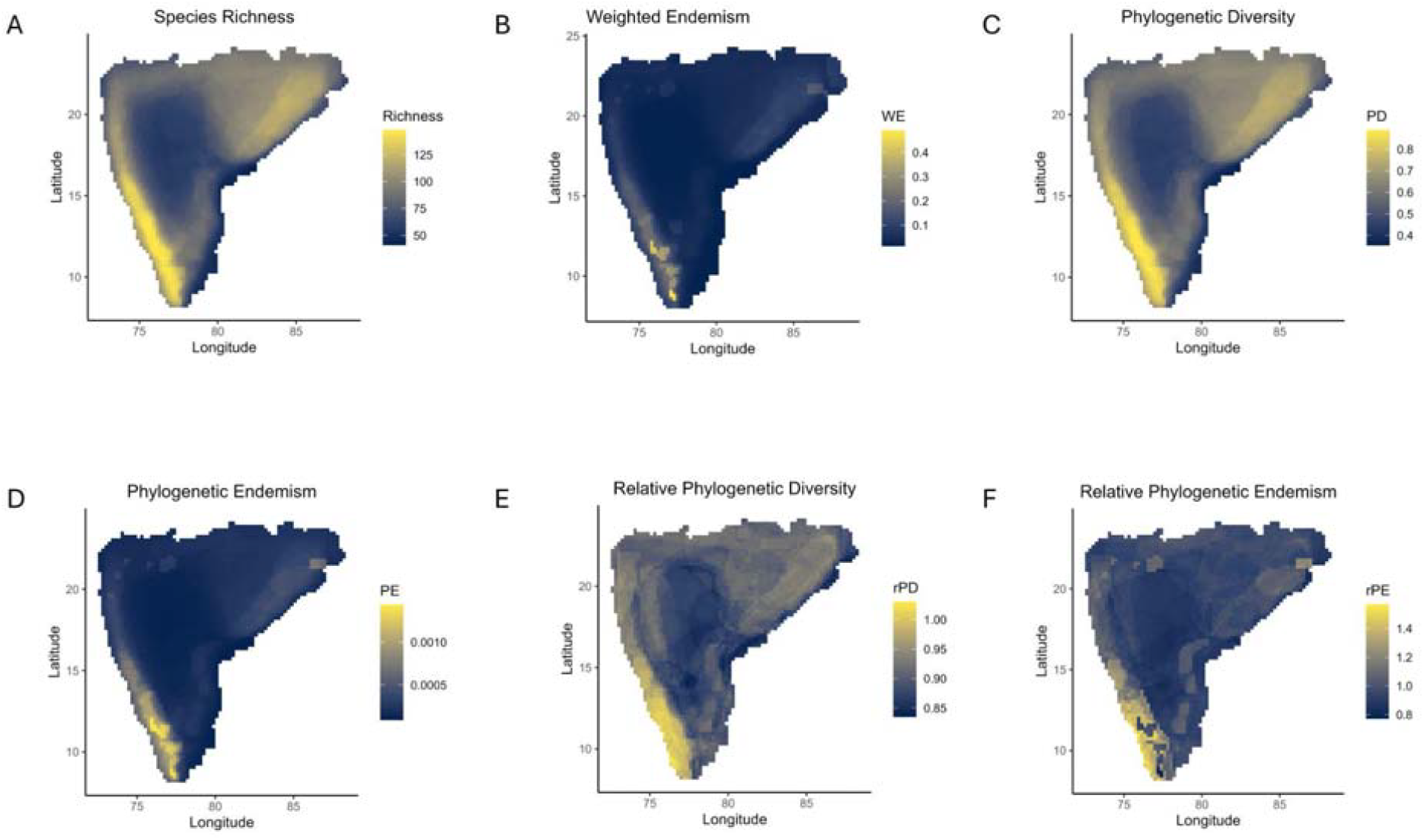
Spatial patterns of forest bird diversity across peninsular India at a 25 km × 25 km resolution based on BirdLife International range maps. (A) Species Richness (SR), (B) Weighted Endemism (WE), (C) Phylogenetic Diversity (PD), (D) Phylogenetic Endemism (PE), (E) Relative Phylogenetic Diversity (RPD), and (F) Relative Phylogenetic Endemism (RPE). The southern Western Ghats exhibit the highest values across all metrics, indicating the presence of ancient, range-restricted lineages. Additional hotspots are observed in parts of the Eastern Ghats and central Indian highlands. The results at all three scales - 50km x 50km, 25km x 25km and 10km x 10km - are mostly concordant (Figures 2, S5, S6 and S7). All the estimates for SR, WE, PD, PE, RPD, RPE, and TILD are consistent with highest diversity in the southern WG compared to the rest of the peninsula. Moreover, with eBird data at 25km x 25km scale the patterns in SR, WE, PD, PE are similar to results with birdlife range maps, however there are several gaps for our species which may yield underestimated ranges (Figure S8). Hence, we focus our discussion based on birdlife range maps at a scale of 25km x 25km.

Patterns in Phylogenetic Diversity (PD) also show similar patterns to SR (Figure 2C). The WG shows a latitudinal gradient in PD with increasing PD at lower latitudes. The hill ranges like Satpuras, Chota Nagpur plateau, and the Eastern Ghats also show relatively high PD compared to parts of the rest of peninsular India. Linear regression examining relationships between SR and PD shows that they are significantly positively correlated (Figure S4). Patterns in Phylogenetic Endemism (PE) show a latitudinal gradient in Western Ghats, with southern Western Ghats having highest PE in peninsular India (Figure 2D). Several species of forest birds such as Grey-headed Bulbul *Microtarsus priocephalus*, Flame-throated Bulbul *Rubigula gularis*, and two genera of birds, *Sholicola* and *Montecincla*, are endemic to the southern Western Ghats (Robin et al. 2017). These species are evolutionarily distinct and occur in small geographic areas contributing to high PE in southern WG.

We find the highest RPD and RPE values in southern Western Ghats, but other areas like the northern Western Ghats, the Eastern Ghats, the central Indian highlands, and parts of the Satpura hills also show relatively high RPD and RPE (Figure 2E and Figure 2F). The results indicate that areas such as the southern WG, having both high RPD and RPE, have higher accumulation of range restricted ancient lineages of forest birds.

Several sites within the southern WG were identified to be centers of super endemism, paleo endemism and mixed endemism (Figure S9) indicating a significantly high proportion of both range restricted ancient and young lineages of forest birds. Some sites with mixed endemism were also spread across parts of the northern Eastern Ghats, and Satpuras - indicative of a mix of both young and ancient range restricted lineages. However, much of peninsular India had a non-significant endemism (Figure S9).

### Patterns in community structure

We observe distinct forest bird communities across Peninsular India, both in terms of species composition and phylogenetic similarity (Figure 3). At lower values of K, regions such as the Western Ghats (WG), Satpuras, Chota Nagpur Plateau, and Eastern Ghats (EG) segregate from the rest of Peninsular India, highlighting the distinctiveness of hill range communities (Figures S12, S13 and S14). With increasing K, finer-scale partitioning emerges within these ranges. At the optimal clustering value of K = 5 for species turnover (Figure S10) and K = 4 for phylogenetic turnover (Figure S11), the Deccan Peninsula and the eastern coastal plains form a single community, while the hill ranges (WG, EG, Satpuras, and Chota Nagpur Plateau) each form distinct clusters (Figures 3a and 3b). Additionally, we observe turnover within the WG across the Goa Gap and within the EG across the Godavari and Krishna rivers. When analyzing only the turnover component of phylogenetic beta diversity, similar patterns emerge: the WG and other hill ranges segregate at low K values, while at K = 4, the WG forms a single phylogenetically coherent community and the central Indian highlands (Satpuras and Chota Nagpur Plateau) cluster with the northern EG (Figure S14). Turnover across the Goa Gap within the WG becomes evident only at higher clustering levels (K = 8; Figure S14).

**Figure 3.**
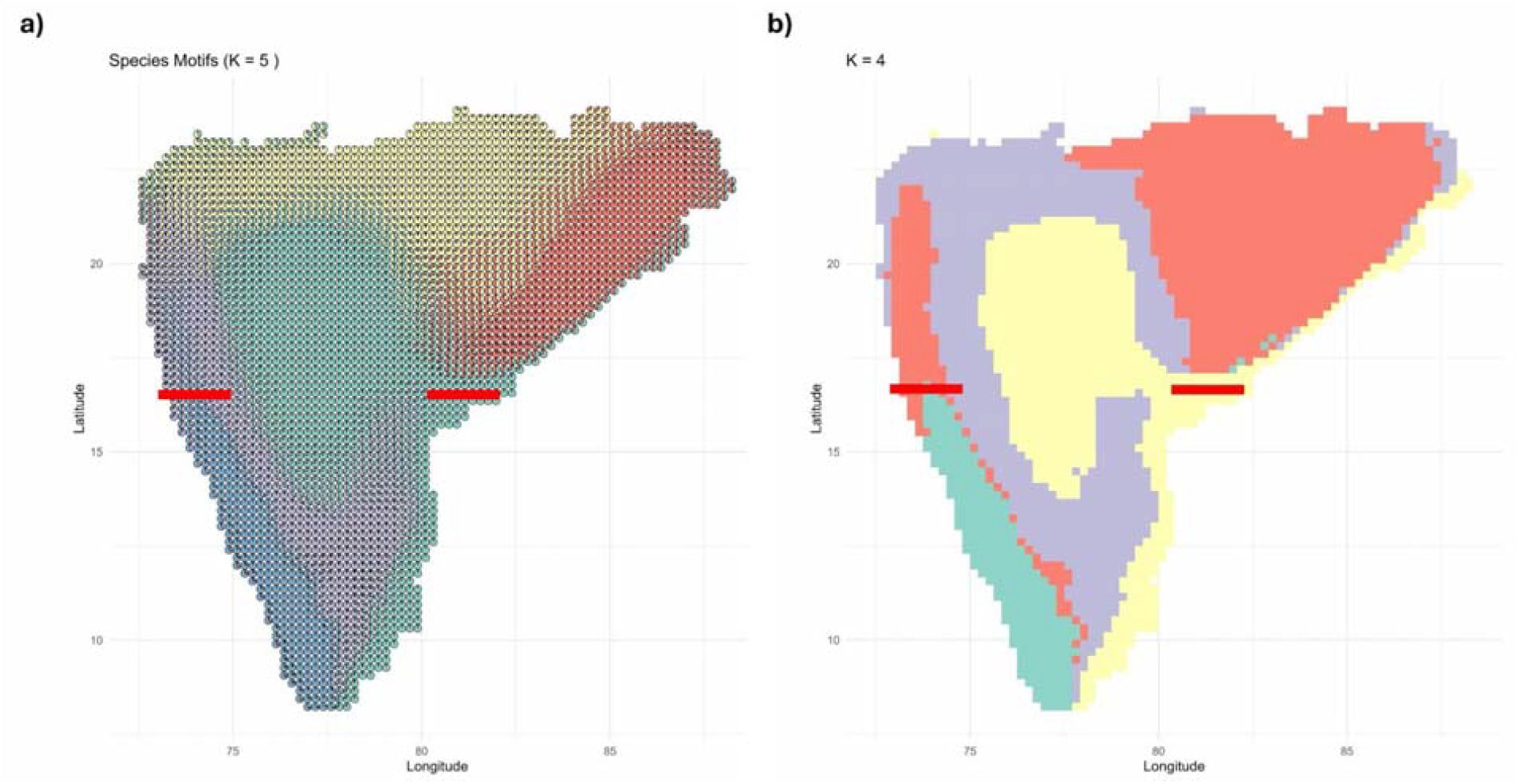
Spatial patterns of forest bird community structure across peninsular India. (A) Taxonomic community structure based on a Grade of Membership model (K = 5), where pie charts show proportions of each community type at each grid cell. (B) Phylogenetic community structure derived from K-means clustering of Sorenson’s phylogenetic beta diversity (K = 4). Distinct clusters correspond to major hill ranges such as the Western Ghats, Satpuras, Chota Nagpur Plateau, and Eastern Ghats, while lowland regions exhibit more homogenous communities. Red lines represent known biogeographic barriers (Ramachandran et al. 2017).

### Predictors of diversity metrics

The best spatial autoregressive models for species richness (SR), phylogenetic diversity (PD), and phylogenetic endemism (PE) included both climate and topographic variables based on AIC scores. Across all three metrics, precipitation of the wettest quarter (Bio16), and topographic ruggedness (TRI) were consistent positive correlates, while climate change velocity since the last glacial maxima (LGM CCV) and mean temperature of the warmest quarter (Bio10) were significant negative correlates (Figure 4).

**Figure 4.**
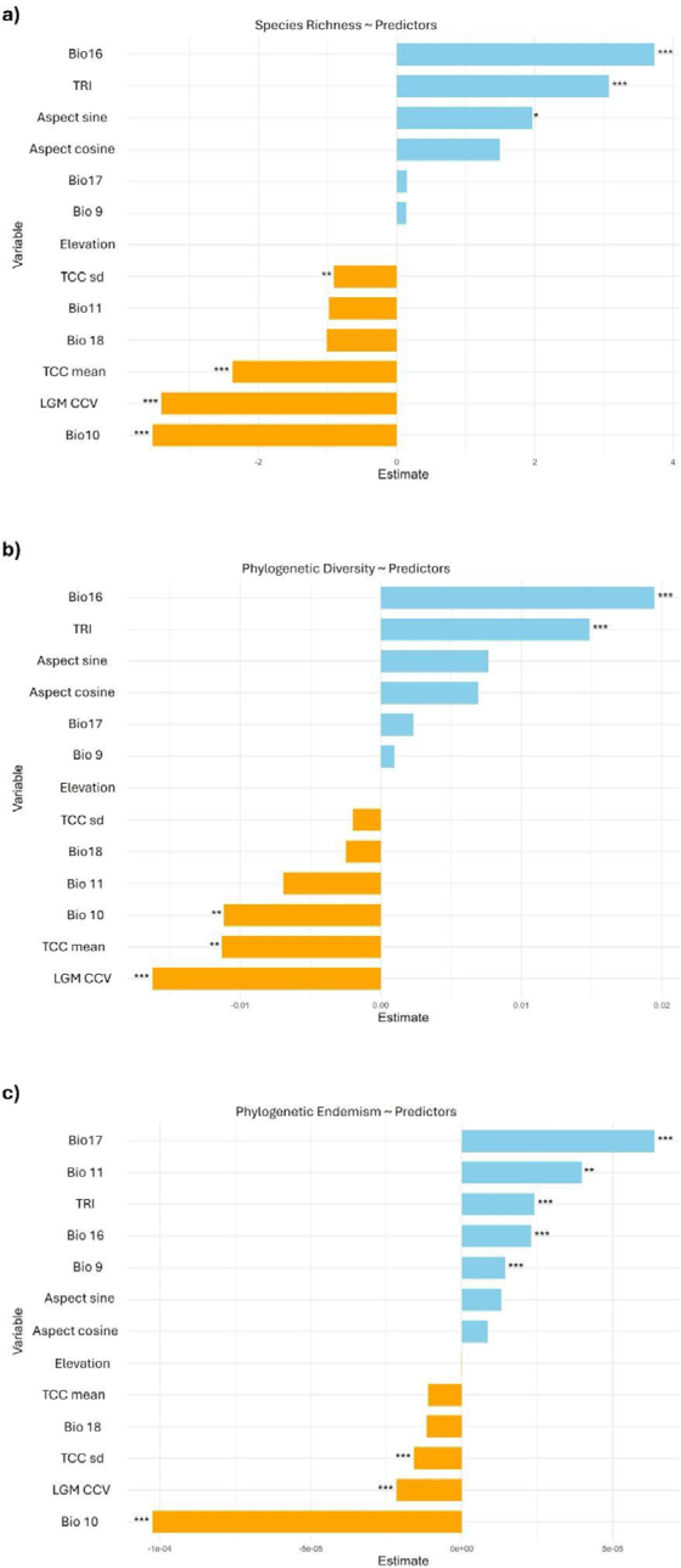
Estimates of coefficients for the best spatial autoregressive models for (a) Species Richness (SR), (b) Phylogenetic Diversity (PD), and (c) Phylogenetic Endemism (PE), incorporating climate and topographic variables. Blue bars indicate positive correlations, and orange bars indicate negative correlations. Significance: *** p < 0.001, ** p < 0.01, * p < 0.05. (a) SR is positively associated with precipitation of the wettest quarter (Bio 16), topographic ruggedness (TRI), and aspect sine, while negatively influenced by mean temperature of the warmest quarter (Bio 10), climate change velocity since LGM (LGM CCV), and total annual cloud cover (TCC mean, TCC sd). (b) PD shows positive associations with Bio 16 and TRI, but negative associations with LGM CCV, Bio 10, and TCC mean. (c) PE is positively correlated with mean temperature of driest quarter (Bio 9), mean temperature of the coldest quarter (Bio 11), precipitation of the driest quarter (Bio 17), Bio 16, and TRI, while Bio 10, LGM CCV, and TCC sd have negative effects.

For SR, additional positive correlates included the sine of aspect, whereas mean annual cloud cover and its seasonality (TCC mean and sd) had negative effects (Figure 4a). The model for PD showed a similar pattern, with mean annual cloud cover having a negative effect (Figure 4b). For PE, precipitation of the driest quarter (Bio17), mean temperature of the coldest quarter (Bio11) were also significant positive correlates. Whereas, the seasonality in annual cloud cover had negative effects on PE (Figure 4c).

## Discussion

Our study quantifies spatial patterns in species richness (SR), weighted endemism (WE), phylogenetic diversity (PD) and phylogenetic endemism (PE) for forest birds of peninsular India. Overall, our results indicate that diversity of forest birds is uneven across peninsular India, with regions like the southern Western Ghats being evolutionary hotspot for forest birds. Moreover, we identify distinct biogeographic zones within the peninsula based on both species composition and phylogenetic similarity. These zones coincide with known biogeographic barriers in highlighting the role of such barriers in shaping the current patterns in bird diversity. We also find support for historic climatic stability and topography playing important roles in driving these patterns in diversity.

### Spatial patterns in diversity of forest birds

The southern Western Ghats (WG) has the highest SR, PD, WE, PE and TILD across the entire peninsula. Further, our results support the idea of latitudinal diversity gradient within the WG with southern WG having higher SR, PD, WE, PE and TILD, decreasing towards higher latitudes. This pattern is known for several taxa in the WG, such as centipedes (Bharti et al. 2021), plants (Gopal et al. 2023; Page and Shanker 2020; Divya, Ramesh, and Karanth 2021; Davidar, Puyravaud, and Leigh 2005), amphibians (Daniels 1992) and snails (Aravind, Rajashekhar, and Madhyastha 2005). Moreover, southern WG also has high relative phylogenetic diversity (RPD) and relative phylogenetic endemism (RPE) for forest birds – indicative of accumulation of several old and range restricted lineages. Similar patterns have been seen in the case of centipedes (Bharti et al. 2021) and plants (Gopal et al. 2023) in the WG with high proportions of old lineages concentrated in the southern WG. Overall, our results support the idea of the southern WG acting as climatic refuge even for birds, allowing the persistence of old lineages over evolutionary timescales.

However, the Eastern Ghats (EG) show a reverse pattern with northern EG having relatively higher SR, WE, PD, PE and TILD than the southern EG. The northern EG and Chota Nagpur Plateau form interesting regions in a biogeographic perspective because of their proximity to the Himalayas. Several Himalayan elements, such as Crimson Sunbird *Aethopyga siparaja*, Lineated Barbet *Psilopogon lineatus*, Grey Treepie *Dendrocitta formosae*, and Abbott’s Babbler *Malacocincla abbotti*, are known to have disjunct distributions within these regions. These elements do not extend further into any other part of the peninsula contributing to the uniqueness of these regions. Furthermore, the northern EG and Chota Nagpur Plateau form a topographically complex region with relatively high rainfall compared to the southern EG due to receiving monsoon from both southwest and northeast monsoon (Ramdas 1974).

Regions like the central Indian highlands including the Satpuras, the Chota Nagpur Plateau, and the northern EG although show relatively high SR, PD and TILD, the WE and PE in these regions are very low compared to regions like the southern WG. Occurrence of very few endemic and range restricted lineages compared to the southern WG may drive these patterns in endemism across the peninsula. These patterns are consistent even for RPD and RPE in these regions. Although RPD is relatively high in northern WG, central Indian highlands, and the northern EG, RPE is low in these regions. This indicates that though there are a high proportion of old lineages in these regions, most of them are widespread and not range restricted. These patterns align with findings from central Indian tropical dry forests, where phylogenetic community analyses of woody trees revealed greater evolutionary distinctiveness in low-elevation plots (Grant et al. 2023).

Categorical analyses of neo and paleo endemism delineated centers of endemism across peninsular India for forest birds. Some sites in the Satpuras, Chota Nagpur Plateau, and the northern EG show mixed endemism type, indicating a mix of both young and old lineages of range restricted species in these regions. While most part of peninsular India has non-significant endemism, the southern WG is a major center of endemism – making it an evolutionary hotspot for forest birds. Areas within southern WG show sites with a mix of both young and old range restricted lineages (mixed and super endemism types), and sites with high accumulation of old range restricted lineages (paleo endemism type). These patterns are consistent with studies on other taxa, such as centipedes (Bharti et al. 2021; Joshi and Karanth 2013) and plants (Gopal et al. 2023), where both old and young range restricted lineages contribute to high diversity in the southern WG.

### Community structure and composition across peninsular India

Our results for taxonomic and phylogenetic turnover in peninsular India identify multiple distinct clusters. We see that the northern EG, and central and southern WG form distinct clusters. Whereas the southern EG and the northern WG show some similarities in their communities. Satpuras and the central Indian highlands form a distinct cluster as well. These patterns are consistent even when we look at the phylogenetic turnover in communities. The turnover is greatest across the Goa gap and the Godavari gap in both taxonomic and phylogenetic turnover. The Godavari gap is a known biogeographic barrier for birds with several sister species split across this barrier (Ramachandran et al. 2017; Ripley and Beehler 1990). Moreover, several Himalayan elements, such as *Aethopyga siparaja, Psilopogon lineatus, Dendrocitta formosae*, and *Malacocincla abbotti*, have their southernmost distribution limited to the north of the Godavari gap. Such species contribute to the uniqueness of the communities of forest birds in the northern EG and central Indian highlands. Similarly, several WG endemics, such as Dark-fronted Babbler *Dumetia atriceps*, Wayanad Laughingthrush *Pterorhinus delesserti*, Grey-headed Bulbul *Microtarsus priocephalus*, and Flame-throated Bulbul *Rubigula gularis*, are known to have their distributional ranges restricted to south of the Goa gap (Ramachandran et al. 2017). Although a previous study on centipedes revealed turnover in communities across the Palghat gap in southern WG (Bharti et al. 2021), we recover no such pattern at any value of K for both taxonomic and phylogenetic turnover in our study. This may be surprising when we consider the fact that Palghat gap is a well-known biogeographic barrier in the southern WG for several taxa (Joshi and Karanth 2013; Biswas and Praveen Karanth 2021), including birds (Robin et al. 2015). However, we believe that this can potentially be an effect of coarseness of the study scale and expert drawn range maps for birds, and investigations at a finer scale may reveal community structure in forest birds across the Palghat gap. Several potential biogeographic units have been identified for birds across peninsular India by investigating turnovers in birds at a smaller taxonomic unit of subspecies (Ramachandran et al. 2017). Our results are mostly concordant with these patterns depicting strongest taxonomic turnover across the Goa gap and Godavari gap in peninsular India. Even in a phylogenetic perspective our results show the effects of Goa gap and Godavari gap in shaping patterns in phylogenetic similarity in forest bird communities in peninsular India, reinforcing the importance of biogeographic barriers in shaping community structure. However, it is important to note that several species from peninsular India do not have genetic datasets available and hence the diversity in the region may be underestimated due to the potential of cryptic species within the region (Reddy 2014).

### Correlates of forest bird diversity in peninsular India

We tested the role of topography and climate in driving patterns in species richness (SR), phylogenetic diversity (PD), and phylogenetic endemism (PE) for forest birds in peninsular India. Our results indicate that both topography and climate play an important role in driving patterns in SR, PD, and PE.

Topographic ruggedness index was found to be a strong and significant positive predictor of all the three diversity metrics for forest birds. Regions with high topographic complexity in peninsular India have high SR,PD and PE. These topographically complex regions coincide with major hills in peninsular India. Global scale studies have shown that mountains tend to harbor high diversity (Rahbek, Borregaard, Antonelli, et al. 2019; Rahbek, Borregaard, Colwell, et al. 2019) compared to nearby lowlands and less topographically complex areas. Mountains with their complex topography have high climatic heterogeneity over small spatial scales and act as barriers to dispersal, ultimately promoting the process of speciation (Rahbek, Borregaard, Antonelli, et al. 2019; Rahbek, Borregaard, Colwell, et al. 2019). The Western Ghats (WG) in peninsular India has high topographical complexity in lower latitudes compared to higher latitudes. Since the complexity of topography was found to be a significant positive predictor for SR, PD and PE, the decrease in diversity in higher latitudes of WG may be the result of reduced topographical complexity in higher latitudes. A similar trend is seen even in the case of plants, where higher evolutionary diversity is seen in topographically heterogenous stretches of the WG (Gopal et al. 2023). The role of topography in driving patterns in SR and PD can be even extended to the Eastern Ghats (EG). Higher SR and PD in northern EG as opposed to southern EG, as expected under predictions from latitudinal diversity gradient, can be explained by heterogeneity in topography across the EG. The northern EG have a higher topographic ruggedness compared to the southern EG, which may potentially drive the pattern of reversed latitudinal diversity gradient in the EG for forest birds. This coupled with geographic proximity of the northern EG to the Himalayas may play a crucial role in persistence of high diversity within the northern EG.

Our results also suggest a major role of climate in driving patterns in SR, PD and PE. Regions with high rainfall (high Bio16, Bio17), moderately warmer climate (high Bio9, Bio11, but low Bio10), and historically stable climate (low quaternary climate change velocity) tend to have higher SR, PD and PE. Our results are in line with expectations that stable, warm and wet climate – climatic refugia - supports higher diversity (Araújo et al. 2008; Carnaval et al. 2009; Gómez et al. 2007; Wiens and Donoghue 2004). Regions like the southern WG are known to have acted as climatic refuge based on the pollen records (Mishra et al. 2024; Prasad et al. 2009) as well as persistence of several ancient lineages within the region (Joshi and Karanth 2011; Gopal et al. 2023; Bharti et al. 2021; Joshi and Karanth 2013; Vijayakumar et al. 2016). Overall climatic stability coupled with complex topography supports the persistence of ancient lineages as well as provides opportunities for younger lineages to emerge in the southern WG – yielding high PD and PE in the region. Our findings are consistent with global patterns across taxa where regions with complex topography and historically stable, warm, and wet climates have been shown to harbor higher species richness, phylogenetic diversity, and endemism (Wiens and Donoghue 2004; Carnaval et al. 2009; Rahbek, Borregaard, Antonelli, et al. 2019; Paúl et al. 2023; Sandel et al. 2011; Fjeldså, Bowie, and Rahbek 2012).

### Caveats

We also note that expert drawn range maps for birds have shown to overestimate the actual range of species distributions, particularly at finer scales (Warudkar et al. 2022). This mismatch of distribution may influence the estimates of diversity at finer spatial scales and caution must be taken in interpreting the results at such scales. This also has an effect on the patterns in endemism recovered as it is known that endemism patterns are scale dependent (Daru et al. 2020). Although our dataset primarily includes forest and woodland dependent birds, some widespread species, particularly in groups like Accipitriformes, Strigiformes, and species such as Rose-ringed Parakeet *Psittacula krameri*, Alexandrine Parakeet *Psittacula eupatria*, and Spotted Owlet *Athene brama*, are difficult to assign to a single habitat type and may introduce noise. Moreover, habitat classifications were sourced from a global trait database (Tobias et al. 2022), which may not accurately represent habitat preferences for species at regional or local scales. Further we note that ∼ 15 % of the species in this study (29 of 188 species) do not have phylogenetic information in the latest birds of the world phylogeny and have been imputed (McTavish et al. 2024). Several of these species are WG endemics (Table S4) which may lead to underestimation of diversity metrics for the WG region - although it may not change much of our current inferences. Lastly, the lack of systematic genetic sampling across peninsular India has been a major limitation in assessing the species limits and uncovering cryptic diversity for several bird species both in a regional and global context (Reddy 2014).

## Conclusion

In summary, our study provides a comprehensive assessment of taxonomic and phylogenetic diversity patterns in forest birds across peninsular India, revealing strong spatial heterogeneity shaped by a combination of historical, climatic, and topographic factors. The southern Western Ghats emerge as a clear evolutionary hotspot, harboring high species richness, endemism, and an accumulation of ancient lineages. Conversely, regions like the northern Eastern Ghats and central Indian highlands (Satpuras and Chota Nagpur Plateau) exhibit unique assemblages influenced by proximity to the Himalayas, and distinct climatic and topography. Our results also underscore the role of biogeographic barriers—particularly the Goa and Godavari gaps—in shaping community structure and turnover. Importantly, we highlight the influence of climatic stability and complex topography in sustaining high diversity and promoting endemism. While our findings are broadly consistent with patterns observed in other taxa, caveats such as missing phylogenetic data, cryptic diversity, and the use of expert drawn range maps underscore the need for finer-scale genetic and distributional data. Taken together, this study adds to our understanding of forest bird diversity in peninsular India and provides critical insights for biogeography in this understudied yet highly biodiverse region and broadly conforms to the global patterns underscoring the importance of historic climatic stability, complex geoclimatic history and topography in driving patterns in diversity.

## Acknowledgements

We acknowledge the Indian Institute of Science Education and Research Tirupati for providing us with computational and infrastructural support. We thank Dr Umesh Srinivasan, Dr Brian Tilston Smith and Dr Rohit Naniwadekar for providing their valuable insights on the study and manuscript. We also thank Aishwarya Bhandari and Vinay KL for their suggestions in improving the manuscript, and Ashwin Warudkar, Chiti Arvind and bird lab members at IISER Tirupati for fruitful discussions and feedback on the study and manuscript. This study was supported by funding from Rohini Nilekani Philanthropies.

## Credit Statement

**Naman Goyal:** Conceptualization (Lead), Data Curation (Lead), Formal Analysis (Lead), Funding Acquisition (Supporting), Investigation (Lead), Methodology (Equal), Project Admin (Equal), Software (Equal), Validation (Equal), Visualization (Lead), Writing Original Draft (Lead), Writing Review & Editing (Equal)

**Archita Sharma:** Funding Acquisition (Supporting), Data Curation (Supporting), Formal Analysis (Supporting), Investigation (Supporting), Methodology (Equal), Software (Equal), Validation (Equal), Visualization (Supporting), Writing Original Draft (Supporting), Writing Review & Editing (Equal)

**Vishwa Jagati:** Data Curation (Supporting), Formal Analysis (Equal), Methodology (Equal), Software (Equal), Validation (Equal)

**Arunima Jain:** Data Curation (Supporting), Formal Analysis (Equal), Methodology (Equal), Software (Equal), Validation (Equal)

**Akshay Herur:** Data Curation (Supporting), Formal Analysis (Equal), Methodology (Equal), Software (Equal), Validation (Equal)

**Abhishek Gopal:** Conceptualization (Supporting), Methodology (Equal), Software (Equal), Validation (Equal), Writing Review & Editing (Equal)

**Jahnavi Joshi:** Conceptualization (Supporting), Methodology (Equal), Supervision (Supporting), Validation (Equal), Writing Review & Editing (Equal)

**VV Robin:**Conceptualization (Supporting), Funding Acquisition (Lead), Investigation (Supporting), Methodology (Equal), Project Admin (Equal), Resources (Lead), Supervision (Lead), Validation (Equal), Writing Original Draft (Supporting), Writing Review & Editing (Equal)

