## SupplementaryMaterial for "Spatial patterns of diversity in forest birds of peninsular India"

**Figure S1** Phylogeny of the 188 species of forest and woodland habitat birds selected in this study

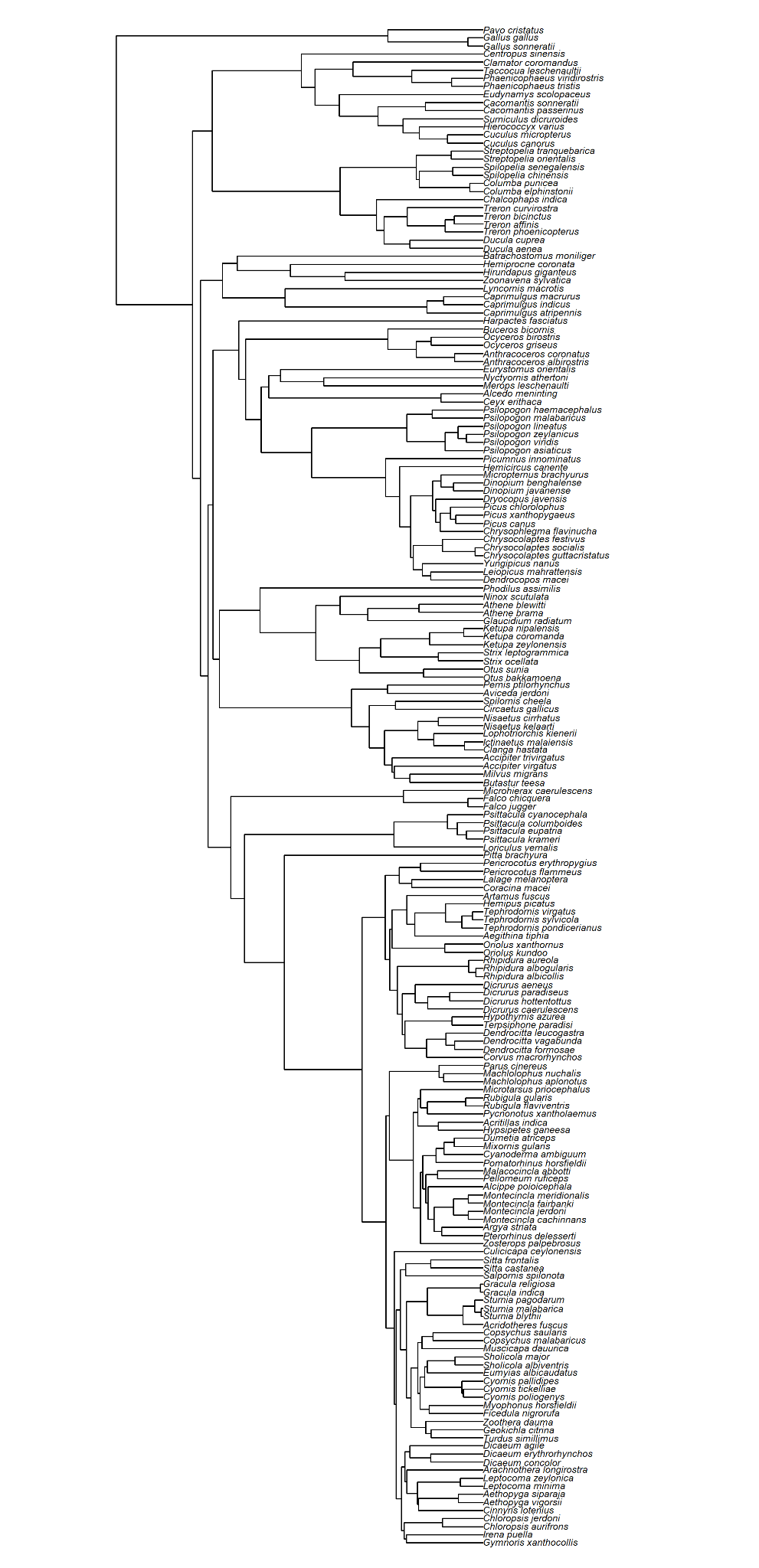

**Figure S2** Optimization for *n_reps* at A) 10km x 10km and B) 25km x 25km scales

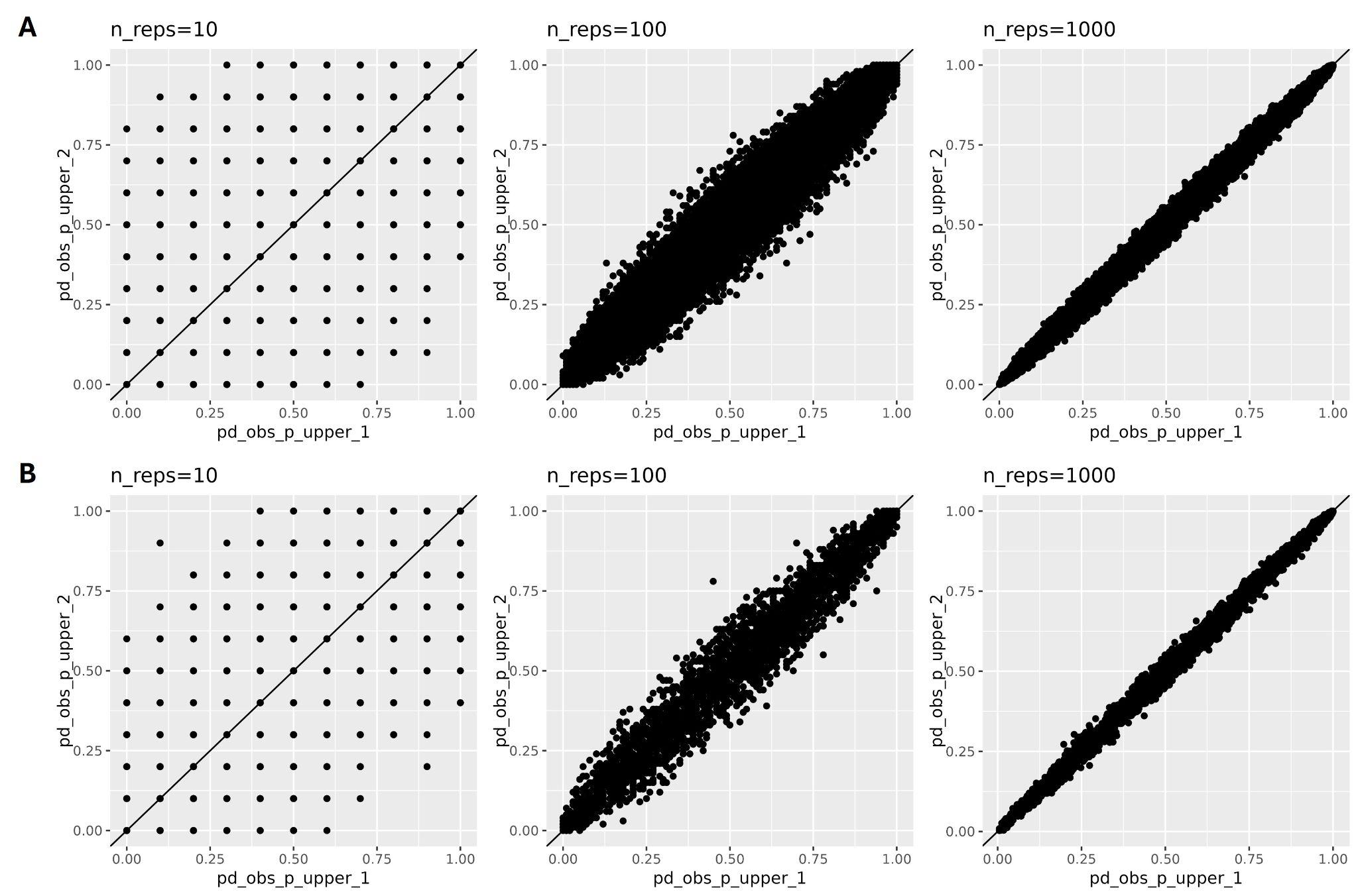

**Figure S3** Optimization for *n_iterations* at A) 10km x 10km and B) 25km x 25km scales

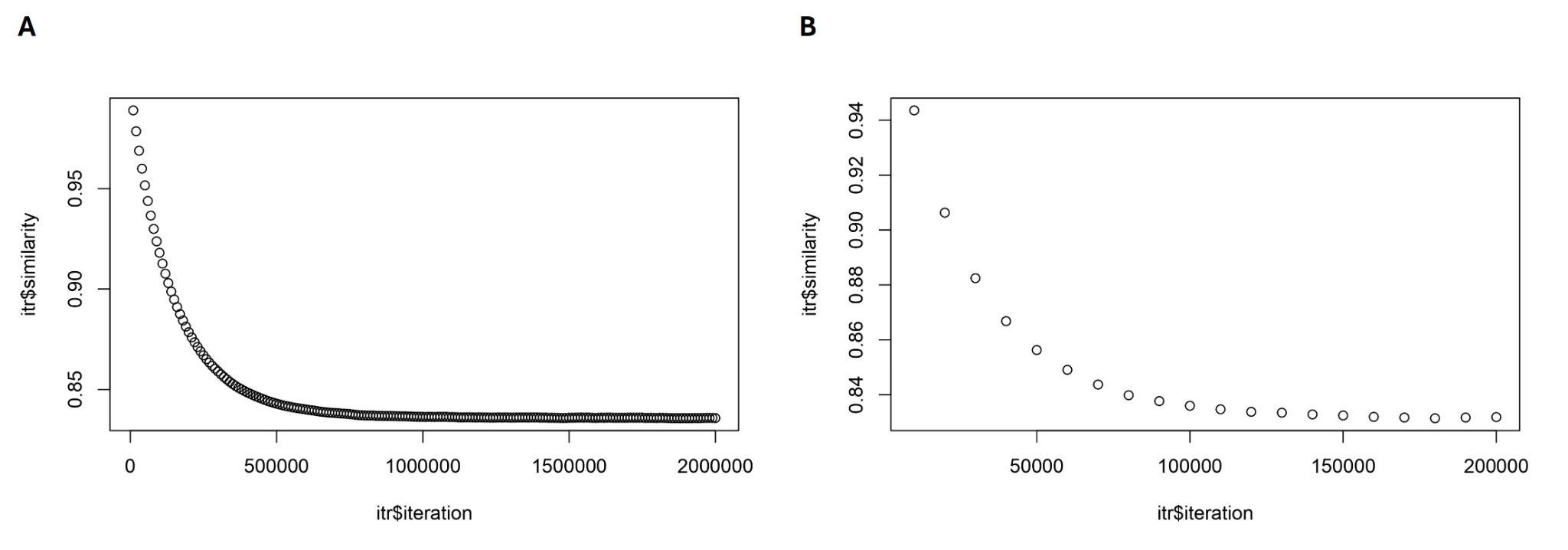

**Figure S4** Linear regression analysis examining relationships between mapped variables (25km x25 km scale for birdlife distribution data). (Abbreviations on the plot - spdv = Species Richness, pd_obs = Phylogenetic Diversity, pe_obs = Phylogenetic Endemism, rpd_obs = Relative Phylogenetic Diversity, and rpe_obs = Relative Phylogenetic Endemism). Linear regression analysis examining relationships between PD, PE, RPD, and RPD show that they are significantly correlated, although there is some scatter in certain relationships.

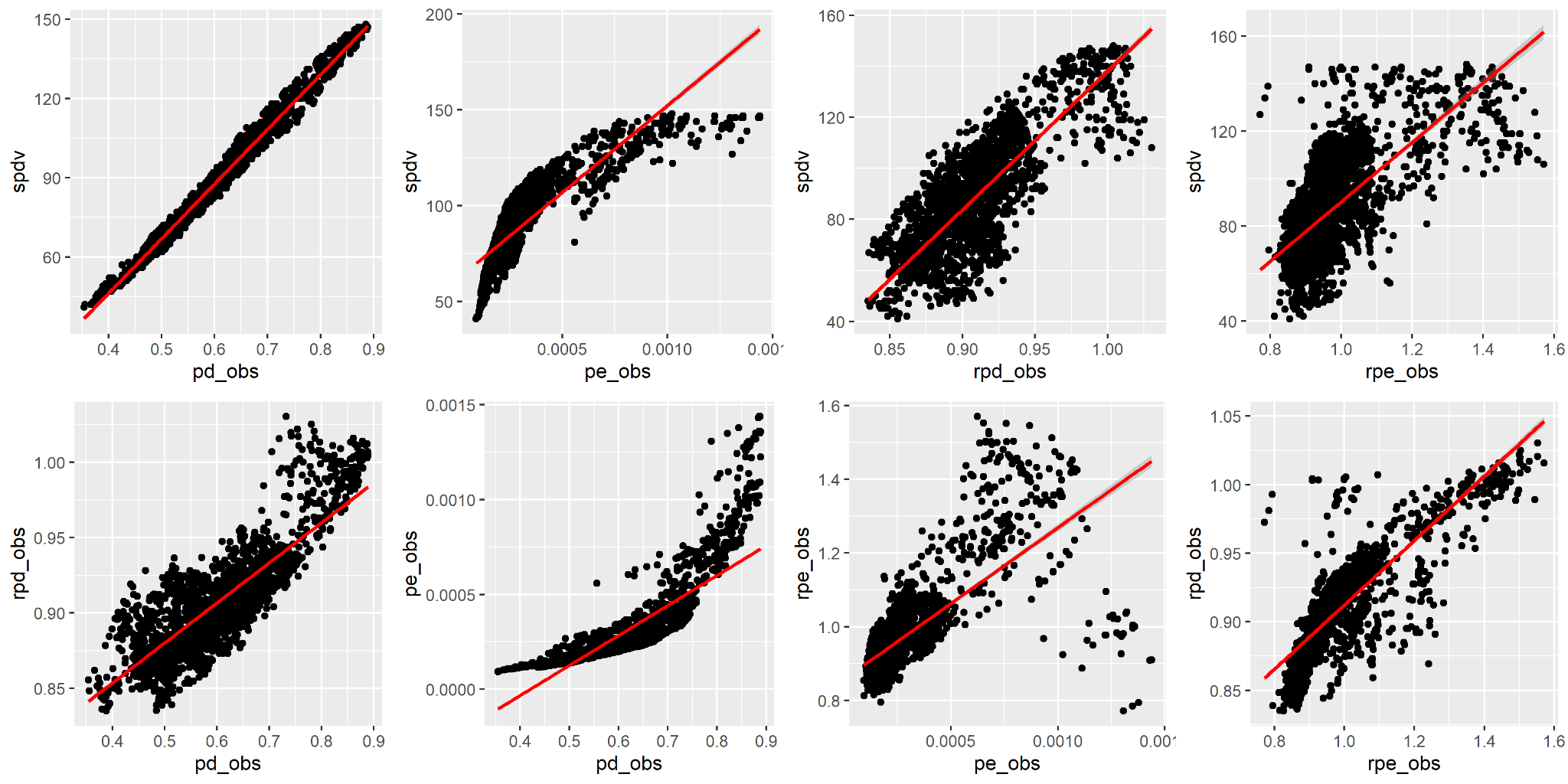

**Figure S5** Patterns in diversity of forest birds in peninsular India at 10x10km scale using BirdLife International range maps, A) Species richness, B) Weighted Endemism (WE), C) Phylogenetic Diversity (PD), D) Phylogenetic Endemism (PE), E) Relative Phylogenetic Diversity (rPD), F) Relative Phylogenetic Endemism (rPE)
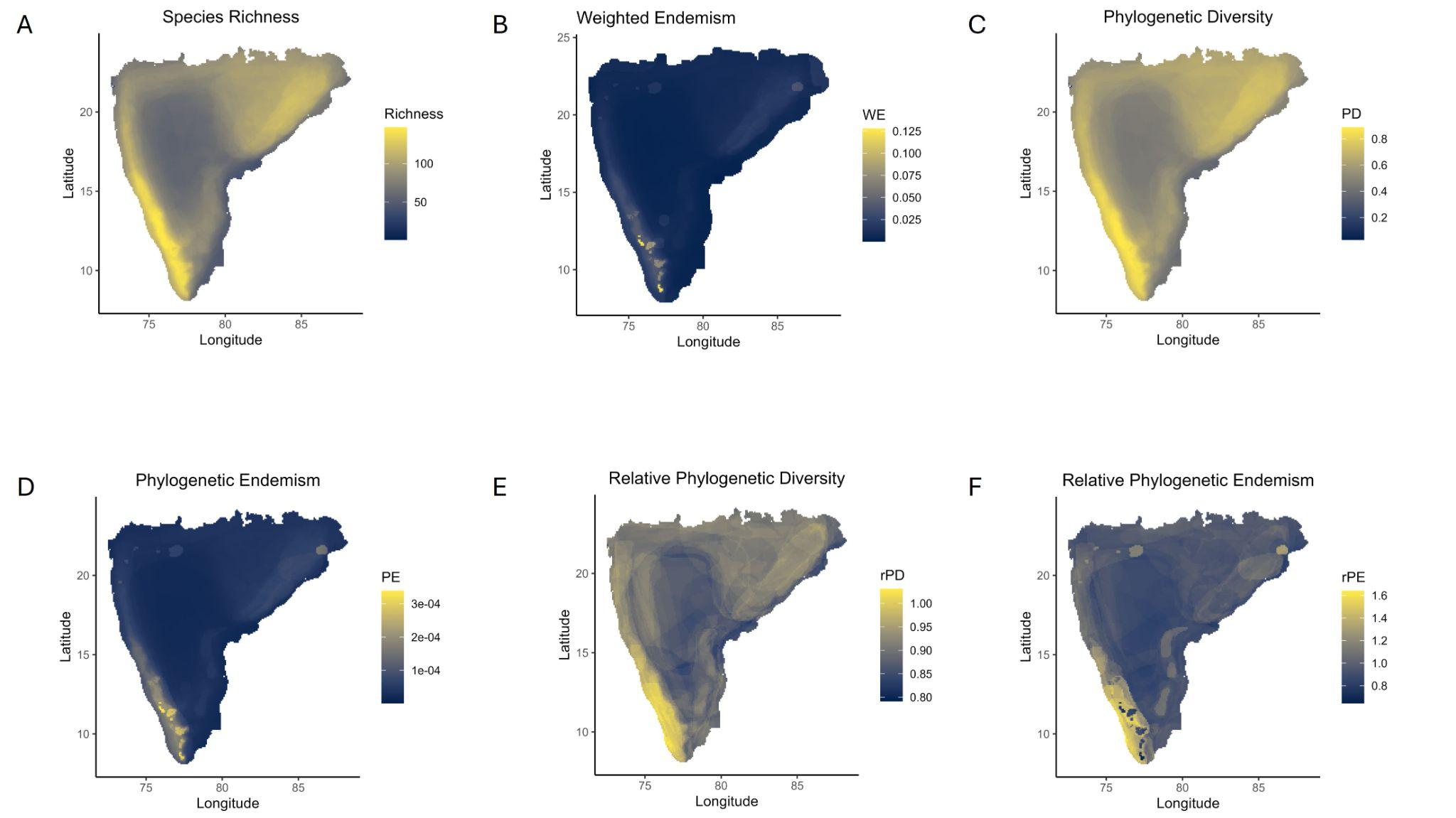

**Figure S6** Patterns in diversity of forest birds in peninsular India at 50x50km scale using BirdLife International range maps, A) Species richness, B) Weighted Endemism (WE), C) Phylogenetic Diversity (PD), D) Phylogenetic Endemism (PE), E) Relative Phylogenetic Diversity (rPD), F) Relative Phylogenetic Endemism (rPE)

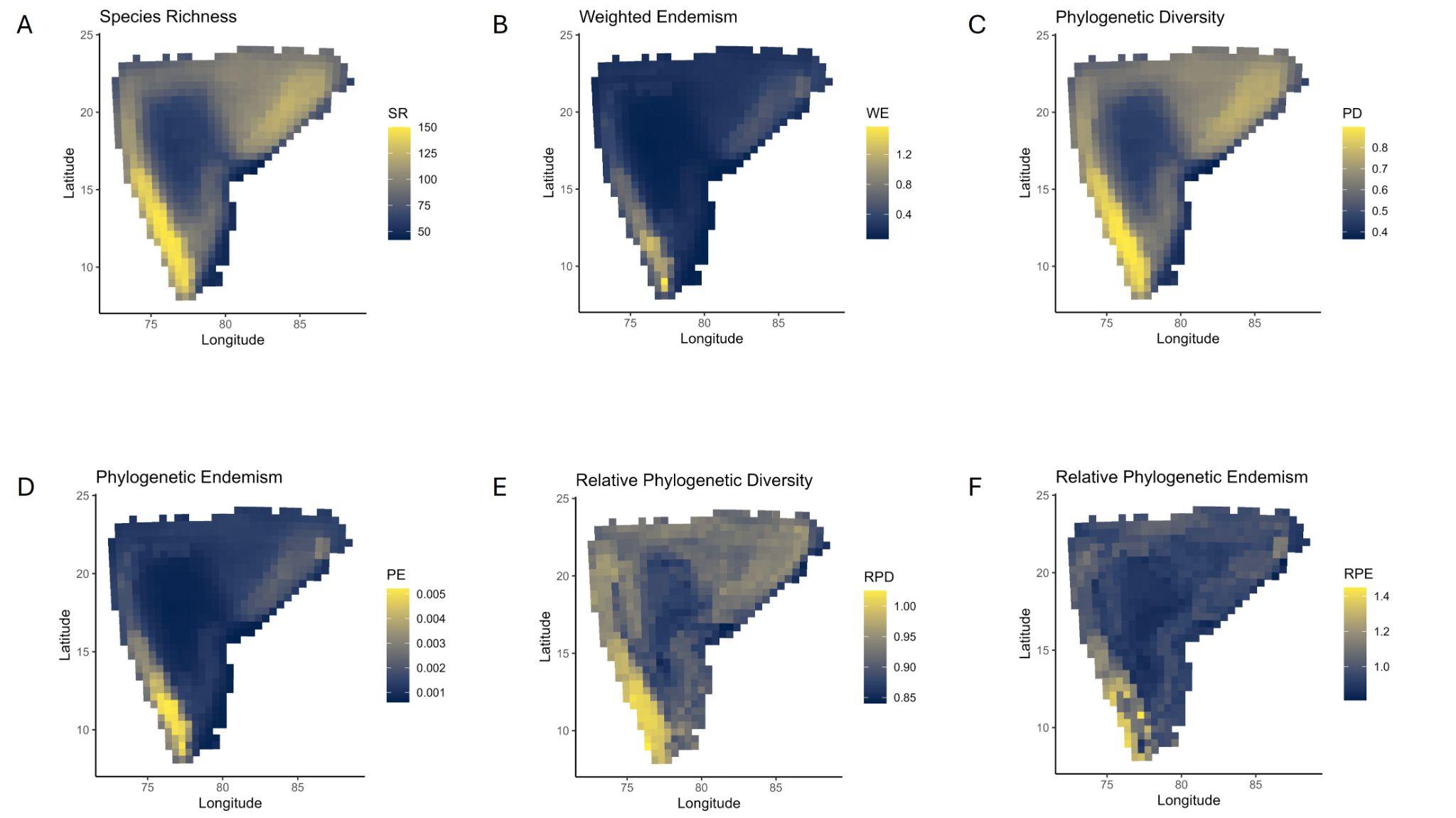

**Figure S7** Time-integrated Lineage Diversity plots for peninsular India at three different scales - A) 50km x 50km, B) 25km x 25km, C) 10km x10km.

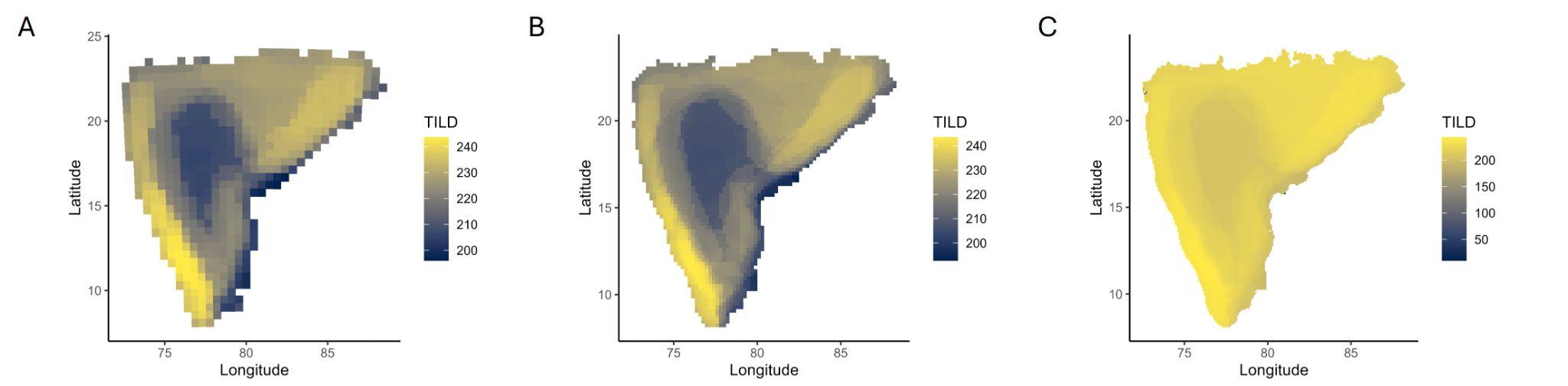

**Figure S8** Patterns in diversity of forest birds in peninsular India at 25x25km scale using eBird data, A) Species richness, B) Phylogenetic Diversity (PD), C) Phylogenetic Endemism (PE), D) Relative Phylogenetic Diversity (rPD), E) Relative Phylogenetic Endemism (rPE), F) Endemism Type. Brighter colours depict areas with high values and darker colours depict areas with low values. Endemism type is classified as Mixed, Neo, Paleo, Super and Non-Signficant based on the method defined in Nitta et al. 2023.

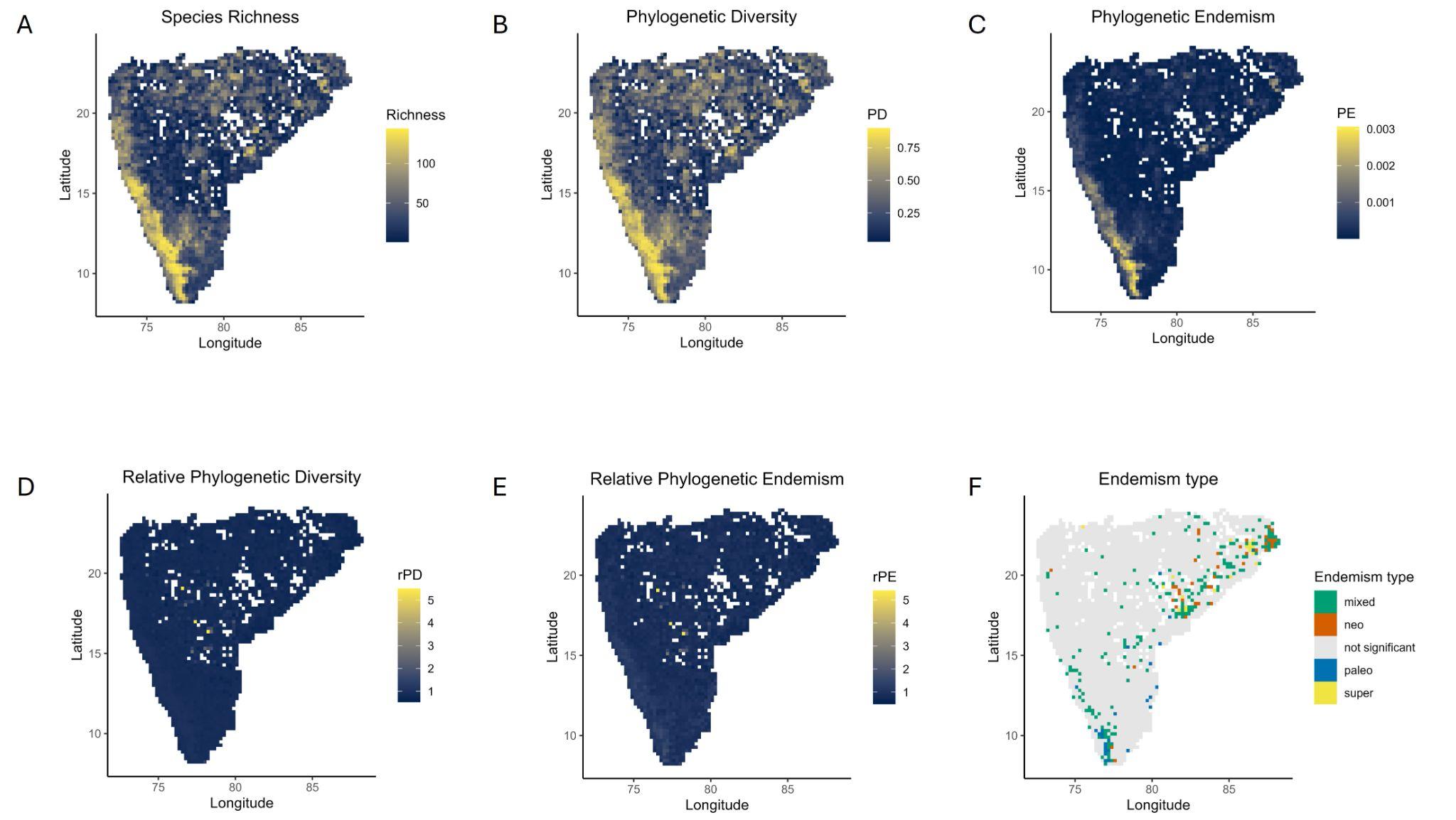

**Figure S9** Endemism Type at three different scales - A) 10km x 10km, B) 25km x 25km, and C) 50km x 50km. Endemism type is classified as Mixed, Neo, Paleo, Super and Non-Signficant based on the method defined in Nitta et al. 2023. Mixed - high proportion of both young and old lineages, Neo - significantly high proportion of young lineages, Paleo - significantly high proportion of old lineages, Super - significantly high proportion of both young and old lineages.

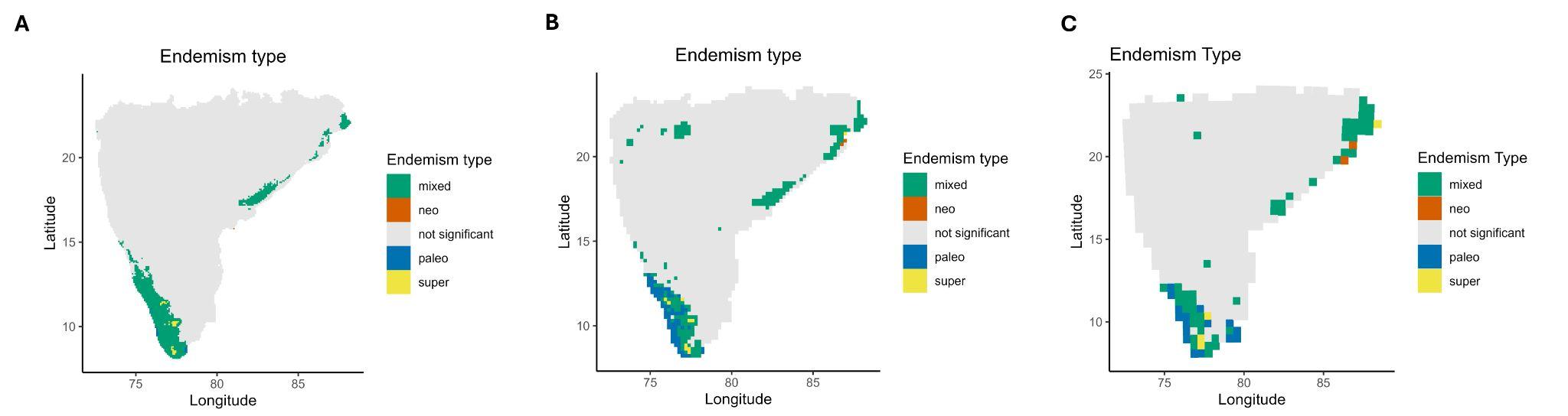

**Figure S10** Elbow plot depicting the optimal value of K at 5 for log likelihood for the best model for each K. The log likelihood curve tends to start flattening out after K at 5.

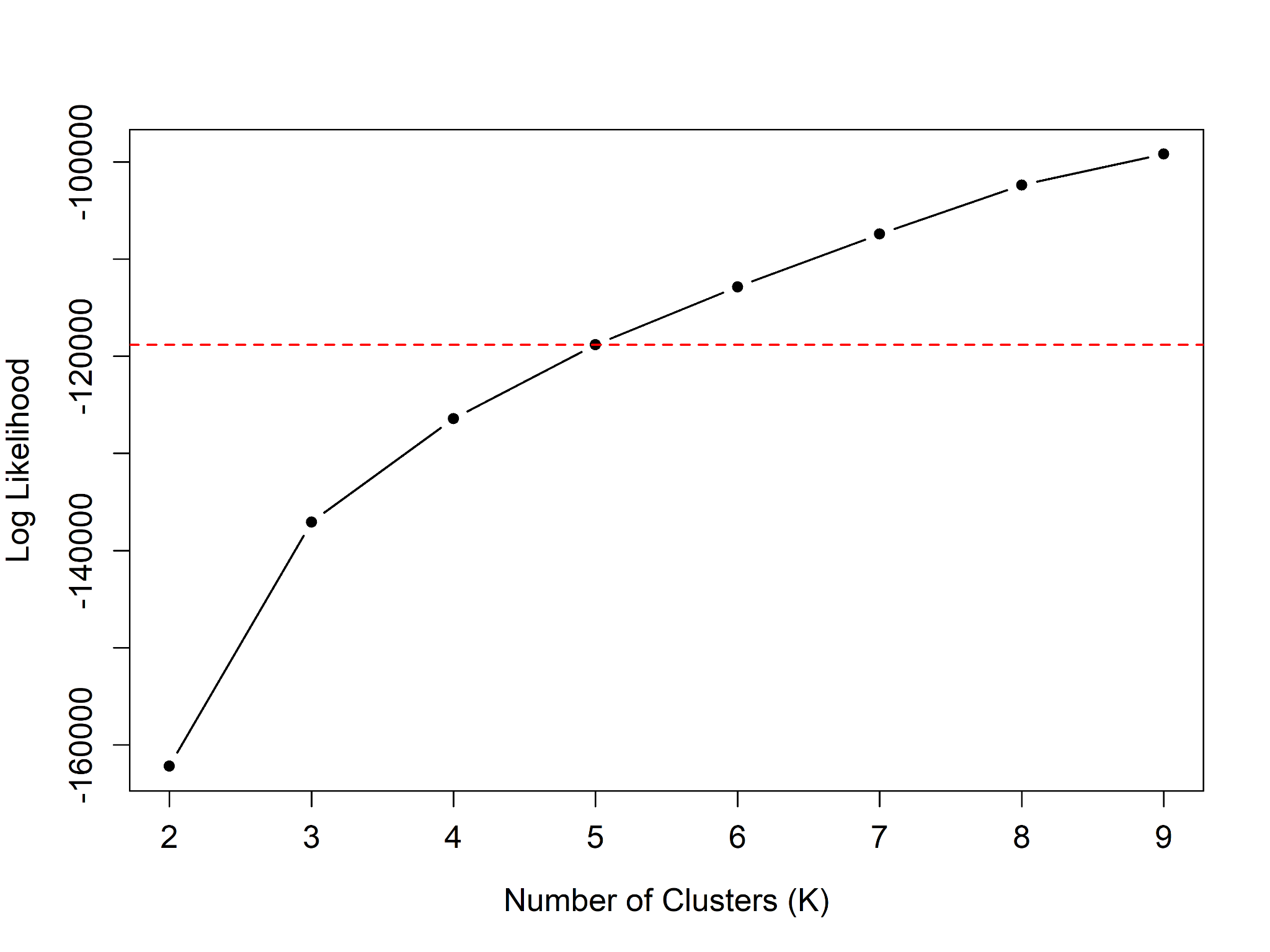

**Figure S11** Elbow plot depicting the best value of K at 4 for K-means clustering of Sorenson’s phylogenetic beta diversity after which the within-cluster sum of squares (WSS) does not decrease significantly.

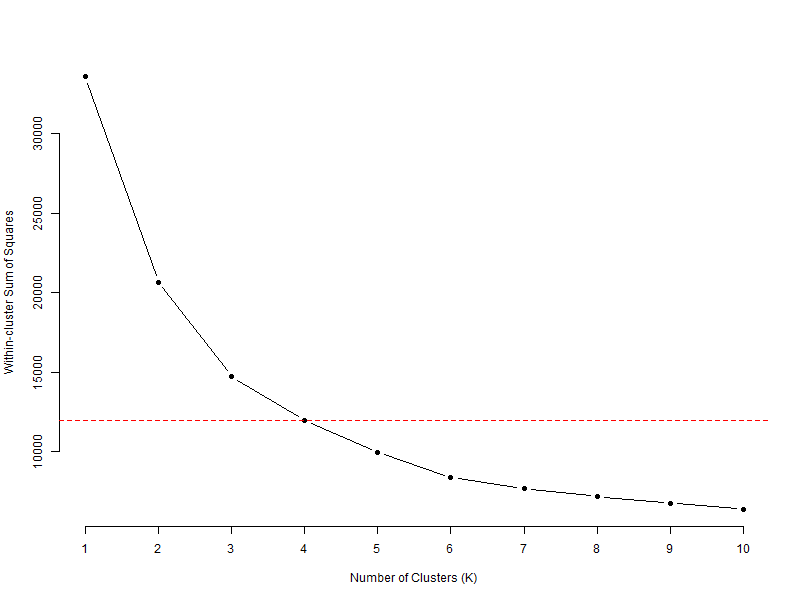

**Figure S12** Patterns in community assemblage of forest birds based on Grade of Membership models fitted on species presence-absence data for values of K ranging from 2 to 9. Here each pie chart on a map represents a grid, and the colours depict the proportion of membership derived from distinct communities based on values of K.

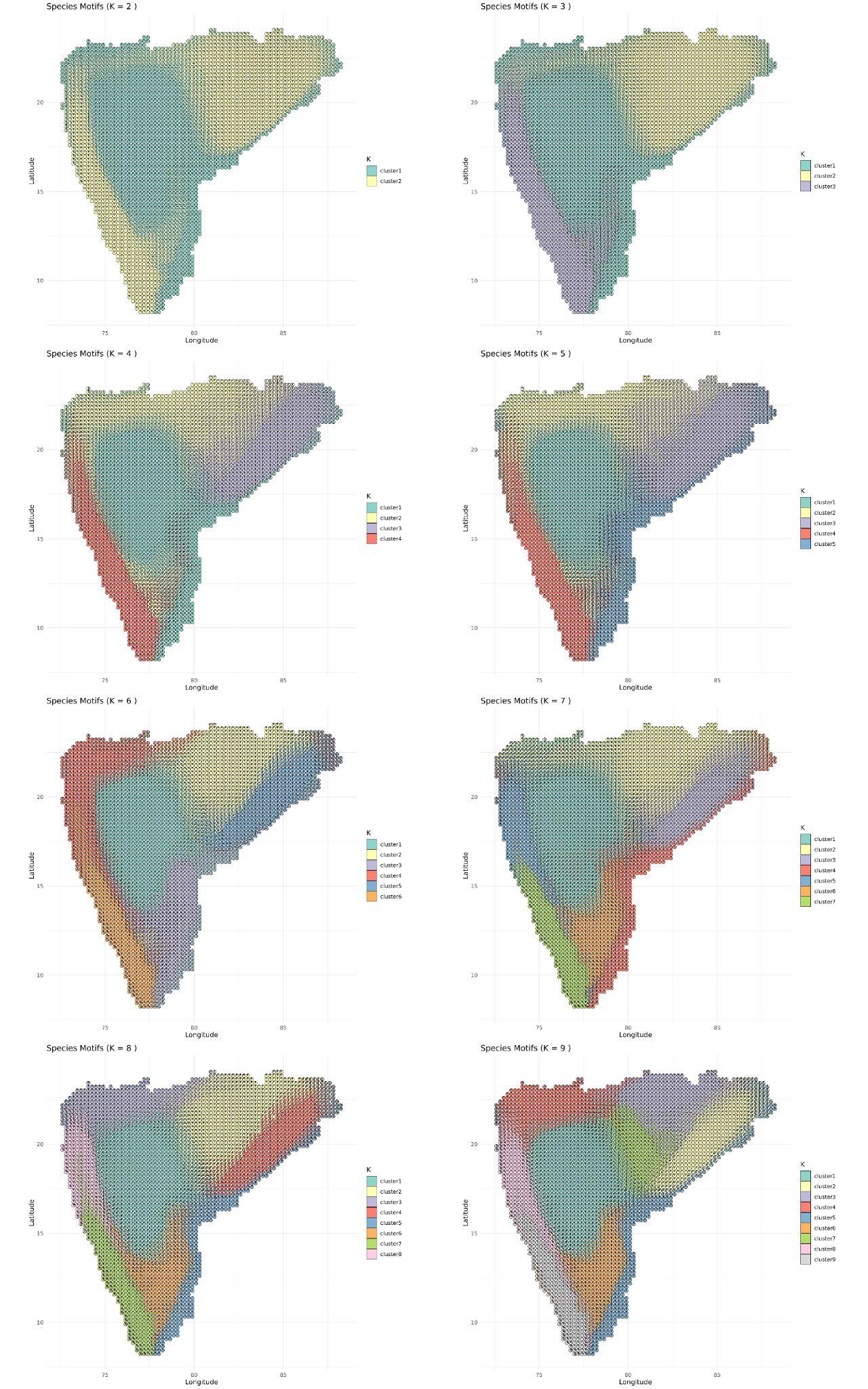

**Figure S13** Patterns in Sorenson’s Phylogenetic beta diversity across peninsular India using K-means clustering algorithm for values of K from 2 to 9. Here each color depicts a cluster of phylogenetically similar regions.

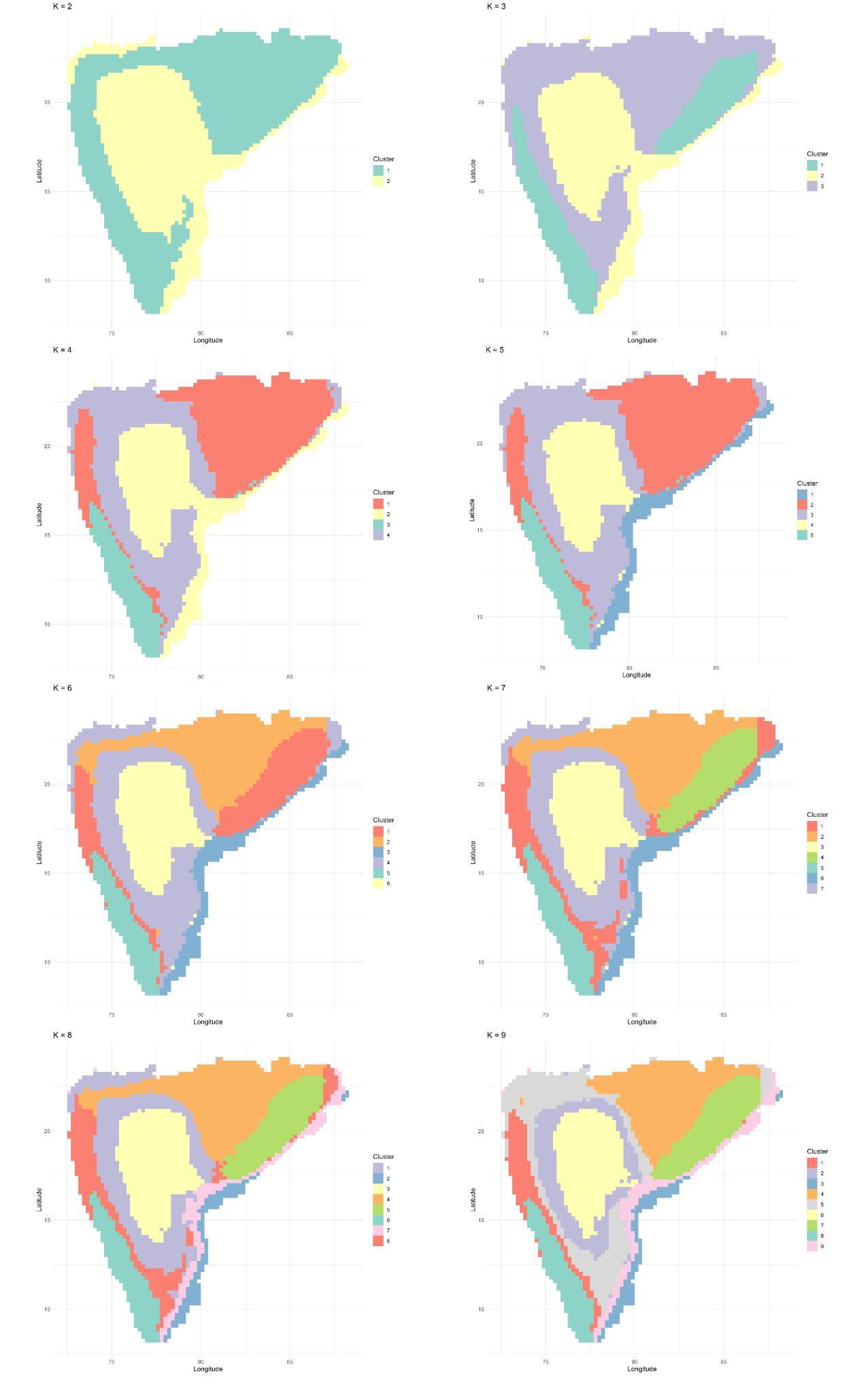

**Figure S14** Patterns in Simpson’s Phylogenetic beta diversity across peninsular India using K-means clustering algorithm for values of K from 2 to 9. Here each color depicts a cluster of phylogenetically similar regions.

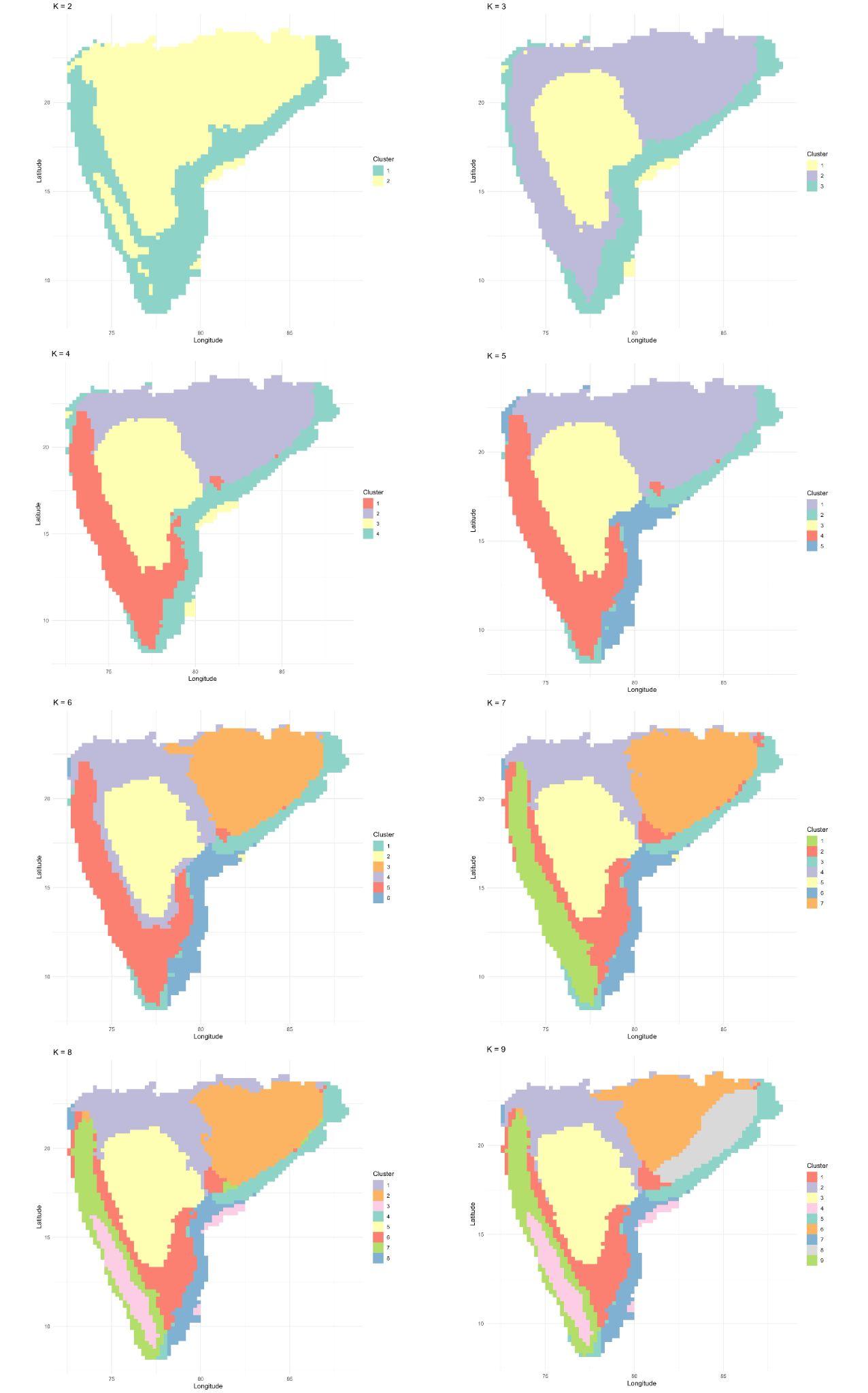

**Table S1** List of resident breeding birds of forest and woodland habitats in peninsular India based on Avonet data (Tobias et al. 2022)

| **Sno** | **Species** | **Order** | **Family** |
| --- | --- | --- | --- |
| 1 | Accipiter trivirgatus | Accipitriformes | Accipitridae (Hawks, Eagles, and Kites) |
| 2 | Accipiter virgatus | Accipitriformes | Accipitridae (Hawks, Eagles, and Kites) |
| 3 | Aviceda jerdoni | Accipitriformes | Accipitridae (Hawks, Eagles, and Kites) |
| 4 | Butastur teesa | Accipitriformes | Accipitridae (Hawks, Eagles, and Kites) |
| 5 | Circaetus gallicus | Accipitriformes | Accipitridae (Hawks, Eagles, and Kites) |
| 6 | Clanga hastata | Accipitriformes | Accipitridae (Hawks, Eagles, and Kites) |
| 7 | Ictinaetus malaiensis | Accipitriformes | Accipitridae (Hawks, Eagles, and Kites) |
| 8 | Lophotriorchis kienerii | Accipitriformes | Accipitridae (Hawks, Eagles, and Kites) |
| 9 | Milvus migrans | Accipitriformes | Accipitridae (Hawks, Eagles, and Kites) |
| 10 | Nisaetus cirrhatus | Accipitriformes | Accipitridae (Hawks, Eagles, and Kites) |
| 11 | Nisaetus kelaarti | Accipitriformes | Accipitridae (Hawks, Eagles, and Kites) |
| 12 | Pernis ptilorhynchus | Accipitriformes | Accipitridae (Hawks, Eagles, and Kites) |
| 13 | Spilornis cheela | Accipitriformes | Accipitridae (Hawks, Eagles, and Kites) |
| 14 | Anthracoceros albirostris | Bucerotiformes | Bucerotidae (Hornbills) |
| 15 | Anthracoceros coronatus | Bucerotiformes | Bucerotidae (Hornbills) |
| 16 | Buceros bicornis | Bucerotiformes | Bucerotidae (Hornbills) |
| 17 | Ocyceros birostris | Bucerotiformes | Bucerotidae (Hornbills) |
| 18 | Ocyceros griseus | Bucerotiformes | Bucerotidae (Hornbills) |
| 19 | Hirundapus giganteus | Caprimulgiformes | Apodidae (Swifts) |
| 20 | Zoonavena sylvatica | Caprimulgiformes | Apodidae (Swifts) |
| 21 | Caprimulgus atripennis | Caprimulgiformes | Caprimulgidae (Nightjars and Allies) |
| 22 | Caprimulgus indicus | Caprimulgiformes | Caprimulgidae (Nightjars and Allies) |
| 23 | Caprimulgus macrurus | Caprimulgiformes | Caprimulgidae (Nightjars and Allies) |
| 24 | Lyncornis macrotis | Caprimulgiformes | Caprimulgidae (Nightjars and Allies) |
| 25 | Hemiprocne coronata | Caprimulgiformes | Hemiprocnidae (Treeswifts) |
| 26 | Batrachostomus moniliger | Caprimulgiformes | Podargidae (Frogmouths) |
| 27 | Chalcophaps indica | Columbiformes | Columbidae (Pigeons and Doves) |
| 28 | Columba elphinstonii | Columbiformes | Columbidae (Pigeons and Doves) |
| 29 | Columba punicea | Columbiformes | Columbidae (Pigeons and Doves) |
| 30 | Ducula aenea | Columbiformes | Columbidae (Pigeons and Doves) |
| 31 | Ducula cuprea | Columbiformes | Columbidae (Pigeons and Doves) |
| 32 | Spilopelia chinensis | Columbiformes | Columbidae (Pigeons and Doves) |
| 33 | Spilopelia senegalensis | Columbiformes | Columbidae (Pigeons and Doves) |
| 34 | Streptopelia orientalis | Columbiformes | Columbidae (Pigeons and Doves) |
| 35 | Streptopelia tranquebarica | Columbiformes | Columbidae (Pigeons and Doves) |
| 36 | Treron affinis | Columbiformes | Columbidae (Pigeons and Doves) |
| 37 | Treron bicinctus | Columbiformes | Columbidae (Pigeons and Doves) |
| 38 | Treron curvirostra | Columbiformes | Columbidae (Pigeons and Doves) |
| 39 | Treron phoenicopterus | Columbiformes | Columbidae (Pigeons and Doves) |
| 40 | Alcedo meninting | Coraciiformes | Alcedinidae (Kingfishers) |
| 41 | Ceyx erithaca | Coraciiformes | Alcedinidae (Kingfishers) |
| 42 | Eurystomus orientalis | Coraciiformes | Coraciidae (Rollers) |
| 43 | Merops leschenaulti | Coraciiformes | Meropidae (Bee-eaters) |
| 44 | Nyctyornis athertoni | Coraciiformes | Meropidae (Bee-eaters) |
| 45 | Cacomantis passerinus | Cuculiformes | Cuculidae (Cuckoos) |
| 46 | Cacomantis sonneratii | Cuculiformes | Cuculidae (Cuckoos) |
| 47 | Centropus sinensis | Cuculiformes | Cuculidae (Cuckoos) |
| 48 | Clamator coromandus | Cuculiformes | Cuculidae (Cuckoos) |
| 49 | Cuculus canorus | Cuculiformes | Cuculidae (Cuckoos) |
| 50 | Cuculus micropterus | Cuculiformes | Cuculidae (Cuckoos) |
| 51 | Eudynamys scolopaceus | Cuculiformes | Cuculidae (Cuckoos) |
| 52 | Hierococcyx varius | Cuculiformes | Cuculidae (Cuckoos) |
| 53 | Phaenicophaeus tristis | Cuculiformes | Cuculidae (Cuckoos) |
| 54 | Phaenicophaeus viridirostris | Cuculiformes | Cuculidae (Cuckoos) |
| 55 | Surniculus dicruroides | Cuculiformes | Cuculidae (Cuckoos) |
| 56 | Taccocua leschenaultii | Cuculiformes | Cuculidae (Cuckoos) |
| 57 | Falco chicquera | Falconiformes | Falconidae (Falcons and Caracaras) |
| 58 | Falco jugger | Falconiformes | Falconidae (Falcons and Caracaras) |
| 59 | Microhierax caerulescens | Falconiformes | Falconidae (Falcons and Caracaras) |
| 60 | Gallus gallus | Galliformes | Phasianidae (Pheasants, Grouse, and Allies) |
| 61 | Gallus sonneratii | Galliformes | Phasianidae (Pheasants, Grouse, and Allies) |
| 62 | Pavo cristatus | Galliformes | Phasianidae (Pheasants, Grouse, and Allies) |
| 63 | Aegithina tiphia | Passeriformes | Aegithinidae (Ioras) |
| 64 | Artamus fuscus | Passeriformes | Artamidae (Woodswallows, Bellmagpies, and Allies) |
| 65 | Coracina macei | Passeriformes | Campephagidae (Cuckooshrikes) |
| 66 | Lalage melanoptera | Passeriformes | Campephagidae (Cuckooshrikes) |
| 67 | Pericrocotus erythropygius | Passeriformes | Campephagidae (Cuckooshrikes) |
| 68 | Pericrocotus flammeus | Passeriformes | Campephagidae (Cuckooshrikes) |
| 69 | Salpornis spilonota | Passeriformes | Certhiidae (Treecreepers) |
| 70 | Chloropsis aurifrons | Passeriformes | Chloropseidae (Leafbirds) |
| 71 | Chloropsis jerdoni | Passeriformes | Chloropseidae (Leafbirds) |
| 72 | Corvus macrorhynchos | Passeriformes | Corvidae (Crows, Jays, and Magpies) |
| 73 | Dendrocitta formosae | Passeriformes | Corvidae (Crows, Jays, and Magpies) |
| 74 | Dendrocitta leucogastra | Passeriformes | Corvidae (Crows, Jays, and Magpies) |
| 75 | Dendrocitta vagabunda | Passeriformes | Corvidae (Crows, Jays, and Magpies) |
| 76 | Dicaeum agile | Passeriformes | Dicaeidae (Flowerpeckers) |
| 77 | Dicaeum concolor | Passeriformes | Dicaeidae (Flowerpeckers) |
| 78 | Dicaeum erythrorhynchos | Passeriformes | Dicaeidae (Flowerpeckers) |
| 79 | Dicrurus aeneus | Passeriformes | Dicruridae (Drongos) |
| 80 | Dicrurus caerulescens | Passeriformes | Dicruridae (Drongos) |
| 81 | Dicrurus hottentottus | Passeriformes | Dicruridae (Drongos) |
| 82 | Dicrurus paradiseus | Passeriformes | Dicruridae (Drongos) |
| 83 | Irena puella | Passeriformes | Irenidae (Fairy-bluebirds) |
| 84 | Alcippe poioicephala | Passeriformes | Leiothrichidae (Laughingthrushes and Allies) |
| 85 | Argya striata | Passeriformes | Leiothrichidae (Laughingthrushes and Allies) |
| 86 | Montecincla cachinnans | Passeriformes | Leiothrichidae (Laughingthrushes and Allies) |
| 87 | Montecincla fairbanki | Passeriformes | Leiothrichidae (Laughingthrushes and Allies) |
| 88 | Montecincla jerdoni | Passeriformes | Leiothrichidae (Laughingthrushes and Allies) |
| 89 | Montecincla meridionalis | Passeriformes | Leiothrichidae (Laughingthrushes and Allies) |
| 90 | Pterorhinus delesserti | Passeriformes | Leiothrichidae (Laughingthrushes and Allies) |
| 91 | Hypothymis azurea | Passeriformes | Monarchidae (Monarch Flycatchers) |
| 92 | Terpsiphone paradisi | Passeriformes | Monarchidae (Monarch Flycatchers) |
| 93 | Copsychus malabaricus | Passeriformes | Muscicapidae (Old World Flycatchers) |
| 94 | Copsychus saularis | Passeriformes | Muscicapidae (Old World Flycatchers) |
| 95 | Cyornis pallidipes | Passeriformes | Muscicapidae (Old World Flycatchers) |
| 96 | Cyornis poliogenys | Passeriformes | Muscicapidae (Old World Flycatchers) |
| 97 | Cyornis tickelliae | Passeriformes | Muscicapidae (Old World Flycatchers) |
| 98 | Eumyias albicaudatus | Passeriformes | Muscicapidae (Old World Flycatchers) |
| 99 | Ficedula nigrorufa | Passeriformes | Muscicapidae (Old World Flycatchers) |
| 100 | Muscicapa dauurica | Passeriformes | Muscicapidae (Old World Flycatchers) |
| 101 | Myophonus horsfieldii | Passeriformes | Muscicapidae (Old World Flycatchers) |
| 102 | Sholicola albiventris | Passeriformes | Muscicapidae (Old World Flycatchers) |
| 103 | Sholicola major | Passeriformes | Muscicapidae (Old World Flycatchers) |
| 104 | Aethopyga siparaja | Passeriformes | Nectariniidae (Sunbirds and Spiderhunters) |
| 105 | Aethopyga vigorsii | Passeriformes | Nectariniidae (Sunbirds and Spiderhunters) |
| 106 | Arachnothera longirostra | Passeriformes | Nectariniidae (Sunbirds and Spiderhunters) |
| 107 | Cinnyris lotenius | Passeriformes | Nectariniidae (Sunbirds and Spiderhunters) |
| 108 | Leptocoma minima | Passeriformes | Nectariniidae (Sunbirds and Spiderhunters) |
| 109 | Leptocoma zeylonica | Passeriformes | Nectariniidae (Sunbirds and Spiderhunters) |
| 110 | Oriolus kundoo | Passeriformes | Oriolidae (Old World Orioles) |
| 111 | Oriolus xanthornus | Passeriformes | Oriolidae (Old World Orioles) |
| 112 | Machlolophus aplonotus | Passeriformes | Paridae (Tits, Chickadees, and Titmice) |
| 113 | Machlolophus nuchalis | Passeriformes | Paridae (Tits, Chickadees, and Titmice) |
| 114 | Parus cinereus | Passeriformes | Paridae (Tits, Chickadees, and Titmice) |
| 115 | Gymnoris xanthocollis | Passeriformes | Passeridae (Old World Sparrows) |
| 116 | Malacocincla abbotti | Passeriformes | Pellorneidae (Ground Babblers and Allies) |
| 117 | Pellorneum ruficeps | Passeriformes | Pellorneidae (Ground Babblers and Allies) |
| 118 | Pitta brachyura | Passeriformes | Pittidae (Pittas) |
| 119 | Acritillas indica | Passeriformes | Pycnonotidae (Bulbuls) |
| 120 | Hypsipetes ganeesa | Passeriformes | Pycnonotidae (Bulbuls) |
| 121 | Microtarsus priocephalus | Passeriformes | Pycnonotidae (Bulbuls) |
| 122 | Pycnonotus xantholaemus | Passeriformes | Pycnonotidae (Bulbuls) |
| 123 | Rubigula flaviventris | Passeriformes | Pycnonotidae (Bulbuls) |
| 124 | Rubigula gularis | Passeriformes | Pycnonotidae (Bulbuls) |
| 125 | Rhipidura albicollis | Passeriformes | Rhipiduridae (Fantails) |
| 126 | Rhipidura albogularis | Passeriformes | Rhipiduridae (Fantails) |
| 127 | Rhipidura aureola | Passeriformes | Rhipiduridae (Fantails) |
| 128 | Sitta castanea | Passeriformes | Sittidae (Nuthatches) |
| 129 | Sitta frontalis | Passeriformes | Sittidae (Nuthatches) |
| 130 | Culicicapa ceylonensis | Passeriformes | Stenostiridae (Fairy Flycatchers) |
| 131 | Acridotheres fuscus | Passeriformes | Sturnidae (Starlings) |
| 132 | Gracula indica | Passeriformes | Sturnidae (Starlings) |
| 133 | Gracula religiosa | Passeriformes | Sturnidae (Starlings) |
| 134 | Sturnia blythii | Passeriformes | Sturnidae (Starlings) |
| 135 | Sturnia malabarica | Passeriformes | Sturnidae (Starlings) |
| 136 | Sturnia pagodarum | Passeriformes | Sturnidae (Starlings) |
| 137 | Cyanoderma ambiguum | Passeriformes | Timaliidae (Tree-Babblers, Scimitar-Babblers, and Allies) |
| 138 | Dumetia atriceps | Passeriformes | Timaliidae (Tree-Babblers, Scimitar-Babblers, and Allies) |
| 139 | Mixornis gularis | Passeriformes | Timaliidae (Tree-Babblers, Scimitar-Babblers, and Allies) |
| 140 | Pomatorhinus horsfieldii | Passeriformes | Timaliidae (Tree-Babblers, Scimitar-Babblers, and Allies) |
| 141 | Geokichla citrina | Passeriformes | Turdidae (Thrushes and Allies) |
| 142 | Turdus simillimus | Passeriformes | Turdidae (Thrushes and Allies) |
| 143 | Zoothera dauma | Passeriformes | Turdidae (Thrushes and Allies) |
| 144 | Hemipus picatus | Passeriformes | Vangidae (Vangas, Helmetshrikes, and Allies) |
| 145 | Tephrodornis pondicerianus | Passeriformes | Vangidae (Vangas, Helmetshrikes, and Allies) |
| 146 | Tephrodornis sylvicola | Passeriformes | Vangidae (Vangas, Helmetshrikes, and Allies) |
| 147 | Tephrodornis virgatus | Passeriformes | Vangidae (Vangas, Helmetshrikes, and Allies) |
| 148 | Zosterops palpebrosus | Passeriformes | Zosteropidae (White-eyes, Yuhinas, and Allies) |
| 149 | Psilopogon asiaticus | Piciformes | Megalaimidae (Asian Barbets) |
| 150 | Psilopogon haemacephalus | Piciformes | Megalaimidae (Asian Barbets) |
| 151 | Psilopogon lineatus | Piciformes | Megalaimidae (Asian Barbets) |
| 152 | Psilopogon malabaricus | Piciformes | Megalaimidae (Asian Barbets) |
| 153 | Psilopogon viridis | Piciformes | Megalaimidae (Asian Barbets) |
| 154 | Psilopogon zeylanicus | Piciformes | Megalaimidae (Asian Barbets) |
| 155 | Chrysocolaptes festivus | Piciformes | Picidae (Woodpeckers) |
| 156 | Chrysocolaptes guttacristatus | Piciformes | Picidae (Woodpeckers) |
| 157 | Chrysocolaptes socialis | Piciformes | Picidae (Woodpeckers) |
| 158 | Chrysophlegma flavinucha | Piciformes | Picidae (Woodpeckers) |
| 159 | Dendrocopos macei | Piciformes | Picidae (Woodpeckers) |
| 160 | Dinopium benghalense | Piciformes | Picidae (Woodpeckers) |
| 161 | Dinopium javanense | Piciformes | Picidae (Woodpeckers) |
| 162 | Dryocopus javensis | Piciformes | Picidae (Woodpeckers) |
| 163 | Hemicircus canente | Piciformes | Picidae (Woodpeckers) |
| 164 | Leiopicus mahrattensis | Piciformes | Picidae (Woodpeckers) |
| 165 | Micropternus brachyurus | Piciformes | Picidae (Woodpeckers) |
| 166 | Picumnus innominatus | Piciformes | Picidae (Woodpeckers) |
| 167 | Picus canus | Piciformes | Picidae (Woodpeckers) |
| 168 | Picus chlorolophus | Piciformes | Picidae (Woodpeckers) |
| 169 | Picus xanthopygaeus | Piciformes | Picidae (Woodpeckers) |
| 170 | Yungipicus nanus | Piciformes | Picidae (Woodpeckers) |
| 171 | Loriculus vernalis | Psittaciformes | Psittaculidae (Old World Parrots) |
| 172 | Psittacula columboides | Psittaciformes | Psittaculidae (Old World Parrots) |
| 173 | Psittacula cyanocephala | Psittaciformes | Psittaculidae (Old World Parrots) |
| 174 | Psittacula eupatria | Psittaciformes | Psittaculidae (Old World Parrots) |
| 175 | Psittacula krameri | Psittaciformes | Psittaculidae (Old World Parrots) |
| 176 | Athene blewitti | Strigiformes | Strigidae (Owls) |
| 177 | Athene brama | Strigiformes | Strigidae (Owls) |
| 178 | Glaucidium radiatum | Strigiformes | Strigidae (Owls) |
| 179 | Ketupa coromanda | Strigiformes | Strigidae (Owls) |
| 180 | Ketupa nipalensis | Strigiformes | Strigidae (Owls) |
| 181 | Ketupa zeylonensis | Strigiformes | Strigidae (Owls) |
| 182 | Ninox scutulata | Strigiformes | Strigidae (Owls) |
| 183 | Otus bakkamoena | Strigiformes | Strigidae (Owls) |
| 184 | Otus sunia | Strigiformes | Strigidae (Owls) |
| 185 | Strix leptogrammica | Strigiformes | Strigidae (Owls) |
| 186 | Strix ocellata | Strigiformes | Strigidae (Owls) |
| 187 | Phodilus assimilis | Strigiformes | Tytonidae (Barn-Owls) |
| 188 | Harpactes fasciatus | Trogoniformes | Trogonidae (Trogons) |

**Table S2** Top species contributing to each cluster at optimal value of K=5 for Ecostructure analyses

| **Cluster1** | **Cluster2** | **Cluster3** | **Cluster4** | **Cluster5** |
| --- | --- | --- | --- | --- |
| Otus bakkamoena | Oriolus kundoo | Cinnyris lotenius | Dendrocopos macei | Gracula indica |
| Parus cinereus | Zoonavena sylvatica | Aethopyga vigorsii | Anthracoceros albirostris | Lyncornis macrotis |
| Centropus sinensis | Salpornis spilonota | Pycnonotus xantholaemus | Caprimulgus macrurus | Ducula cuprea |
| Argya striata | Pitta brachyura | Ninox scutulata | Rhipidura albicollis | Sturnia blythii |
| Spilopelia chinensis | Sitta castanea | Turdus simillimus | Aethopyga siparaja | Rubigula gularis |

**Table S3** Model rankings for Spatial Autoregressive models based on AIC values

| Response variable | Model | AIC |
| --- | --- | --- |
| Species Richness (SR) | Climate+Topography | 23003 |
|  | Climate | 23209.19 |
|  | Topography | 24325.79 |
| Phylogenetic Diversity (PD) | Climate+Topography | -9533.47 |
|  | Climate | -8980.47 |
|  | Topography | -7890.65 |
| Phylogenetic Endemism (PE) | Climate+Topography | -46432.4 |
|  | Climate | -46194.7 |
|  | Topography | -44658.9 |

**Table S4** Species with no phylogenetic information in the McTavish et al. 2024 global bird phylogeny

| **Sno** | **Species** |
| --- | --- |
| 1 | Sturnia blythii |
| 2 | Rhipidura albogularis |
| 3 | Chrysocolaptes socialis |
| 4 | Tephrodornis sylvicola |
| 5 | Columba elphinstonii |
| 6 | Columba punicea |
| 7 | Montecincla meridionalis |
| 8 | Montecincla jerdoni |
| 9 | Ketupa coromanda |
| 10 | Leptocoma zeylonica |
| 11 | Leptocoma minima |
| 12 | Aethopyga vigorsii |
| 13 | Treron affinis |
| 14 | Dendrocitta leucogastra |
| 15 | Machlolophus aplonotus |
| 16 | Chrysocolaptes festivus |
| 17 | Nisaetus kelaarti |
| 18 | Strix ocellata |
| 19 | Sitta castanea |
| 20 | Myophonus horsfieldii |
| 21 | Caprimulgus atripennis |
| 22 | Yungipicus nanus |
| 23 | Athene blewitti |
| 24 | Cinnyris lotenius |
| 25 | Ducula cuprea |
| 26 | Aviceda jerdoni |
| 27 | Zoonavena sylvatica |
| 28 | Hirundapus giganteus |
| 29 | Phodilus assimilis |
